# Unveiling the Functional Connectivity of Astrocytic Networks with AstroNet, a Graph Reconstruction Algorithm Coupled to Image Processing

**DOI:** 10.1101/2024.10.15.618423

**Authors:** L. Zonca, F.C. Bellier, G. Milior, P. Aymard, J. Visser, A. Rancillac, N. Rouach, D. Holcman

## Abstract

Astrocytes form extended intercellular networks, displaying complex calcium activity. However, the specific organization of these astrocytic networks and the precise extent of their functional connectivity in different brain areas remain unexplored. To unveil the functional architecture of astrocytic networks, we developed, using a data-driven methodology, a novel algorithm called AstroNet that uses two-photon calcium imaging to map temporal correlations in activation events among neighboring astrocytes. Our approach involves reconstructing functional astrocytic networks by organizing individual astrocyte activation events chronologically. This chronological order creates activity paths that enable the extraction of local astrocyte functional correlations. Ultimately, by tallying the occurrences of direct co-activations between pairs of cells along these pathways, we construct a graph that mirrors the underlying astrocyte functional network. By applying this method to two distinct brain regions (CA1 hippocampus and motor cortex), we identified notable differences in local network organizations in sub-regions of around 20-40 astrocytes. Specifically, the cortex exhibited a lower connectivity, while astrocytes in the hippocampus displayed stronger connections. Moreover, we found that in both regions, astrocytic networks consist of smaller, tightly connected sub-networks embedded within a larger, more loosely connected one. Altogether, our innovative method enables the identification of activation paths among astrocytes, facilitates the characterization of local network functional connectivity, and quantifies distinct connectivity patterns among astrocytes from different brain regions. This approach sheds light on the heterogeneous functional organization of astrocytic networks within the brain, pointing to region-specific astrocyte connectivity.

## Introduction

Glial cells form intricate networks interconnected by gap junctions [1, 2], facilitating the exchange of ions and small molecules. Among these glial cells, astrocytes play a crucial role in various functions [3, 4], including the regulation of potassium flows during neuronal activity [5, 6], ensuring metabolic needs, influencing the formation of synapses, and modulating neuronal circuit activity [7, 8]. At the morphological level, astrocytes establish contact with hundreds of neurons and interact with hundreds (in cortical columns) to thousands (in the hippocampus) of other astrocytes [1]. Despite this extensive structural connectivity, little is known about the motifs of astrocyte calcium signaling at the network level and their specific paths during spontaneous activity.

In neuronal networks, signal transmission occurs through chemical synapses, initiating a chain of events leading to the propagation of action potentials. However, in astroglial networks, only a subset of astrocytes are activated following a calcium burst initiated in an astrocyte [9, 10, 8, 11, 12]. Thus the principles governing the extent and synchronization of calcium activity within astroglial networks remain unclear. We here posit that the activated network’s path could serve as a basis for reconstructing synchronized astrocytes belonging to a preferentially connected network. This hypothesis provides a potential avenue for understanding the selective activation patterns in astrocytic networks during various physiological processes.

Calcium imaging analysis has unveiled recurrent activations within neuronal networks [13], including Up-Down state activity [14]. These networks can range in size, from a few cells in cultured environments [13], to extensive populations exceeding millions in brain slices or in-vivo settings [15]. In contrast, much less is known about such recurrent activity in astrocytes, particularly regarding the local connectivity patterns. Reconstructing connectivity in dissociated cultured neurons has leveraged spike dynamics sorting during spontaneous activity and numerical simulations [16]. Techniques like constructing correlation matrices for place cells in maze-running mice [17], or using calcium fluorescence imaging with statistical inference to discern precise spike timings between neurons and reconstruct networks [18], have been successful in the neuronal domain. Meanwhile, studies have tracked individual neurons across multiple sessions using spatial and temporal metrics [19]. Calcium bursting events in neurons and astrocytes can now be simulated effectively [20], and recent advances in image segmentation have disentangled both astrocytic and neurotransmitter fluorescence dynamics at single-cell and population levels [21]. While a segmentation method has been developed to study calcium bursting events in astrocytes [20], it falls short in extracting causal relationships between neighboring cells and reconstructing local network connectivity. Astrocytic connectivity plays a key role in shaping calcium wave signaling and propagation across networks, as highlighted by [12]. Given that astrocytes can stabilize neuronal firing rates through calcium waves and direct regulation of ionic concentrations, high levels of astrocytic connectivity may help prevent excessive neuronal synchronization, as suggested by [22]. Notably, three-dimensional simulations of astrocyte networks by [23] revealed the existence of astrocyte hubs, which are connected to over 75% of the network, underscoring the potential for astrocytes to act as central regulators within the brain’s signaling architecture. In this study, we analyzed the spontaneous activation patterns exhibited by astrocytes, as revealed by calcium fluorescent imaging using GCaMP6f in acute slices of mice. We present here a computational approach to reconstruct Astrocyte networks and extract connectivity properties from correlated calcium events between Regions of Interest (ROI). With a microscopy field of view (40x), we evaluate a region of 150 ∗ 150*μm*^2^, showing tens of visible astrocytes. This sample is in principle sufficiently to reflect broader astrocytes network activity. Additionally, this approach could be used to reconstruct astrocytic networks from calcium events recorded in-vivo using for example miniscope (one-photon) technology. Through the development of a computational method and algorithm called AstroNet, we used time series data from two-photon calcium imaging of astrocytes to reconstruct local network organizations (see Fig.1). Our analysis involves examining calcium activity in dozens of astrocytes across various recording sessions spanning a few minutes. Following the segmentation and time ordering of astrocytic calcium bursts, we mapped the temporal causal correlation between successively activated astrocytes into paths, revealing levels of connectivity. This critical step enabled the construction of a local astrocytic network structure represented in a graph. We applied this method to recover astrocytic connectivities in the hippocampus (CA1 area) and the motor cortex (CTX). Finally, we derived various statistics associated with these graphs, such as the number of connected astrocytes, the count of highly connected astrocytes (hubs), and the properties of the connectivity matrices of these two regions, reflecting local functional connectivity.

## 1 Results

How astrocytes organize and communicate in network remains unclear, we thus developed a computational approach to reconstruct the local astrocyte functional network based on the assumption that two consecutive calcium events occurring in two neighboring astrocytes reflect a local interaction or communication. There are few calcium events at the same time, as such events last several hundreds of milliseconds, and the size of the network is of the order of tens of astrocytes maximum in the field of view. This direct activation is a form of communication, possibly through gap junctions, relying on calcium-induced-calcium-release process [12]. The overall reconstruction of the astrocytic network relies on this assumption. Thus to reconstruct the local astrocyte network functional connectivity, we first identified the position of active astrocytes displaying calcium signaling detected in brain slices via the selective expression of the GCaMP6 genetically-encoded calcium indicator. Given that GCaMP expression may leave only a very small fraction of astrocytes unlabeled, it is difficult to evaluate here the potential consequences of invisible (unlabeled) astrocytes. However, this should not influence significantly the network statistics we extracted here. In addition to unlabeled astrocytes in the network, there could be ROIs for which the calcium fluorescent signal never passes the activation threshold (see Methods section 3.3.2). These ROIs are discarded and we consider that there are not significant contributors to the spread of calcium signals within the network. This hypothesis is at the basis of the present reconstruction method.

From each active astrocyte ROI, we extracted the individual fluorescence time-series. We then segmented these time-series into calcium bursting events. A calcium event in a ROI (method section 3.3), is counted here when the amplitude exceeds the standard deviation of the completely inactive periods computed along the calcium fluorescence signal, after we removed a moving baseline. The results are segmented events that last few seconds (fig. S2). This pipeline (Fig. 1) is used to reveal the unknown underlying network dynamics (Fig. 1A-B). Briefly, the ROIs are detected by averaging over time light intensity of the whole recording session and then finding the active regions of the image, corresponding to the most luminous ones (Fig. 1C1-C2 and Methods). We then extract the average calcium time-series for each ROI and segment them to detect the activation events by extracting local calcium peaks (Fig. 1C3-C5). To define sequences of activation, that we call co-activation paths, we segment the recording session into *N* subperiods (Fig. 1D1) and identify the active ROIs within each subsection. To extract the co-activations paths (Fig. 1D2), we order the individual activation peaks in time, to construct a sequence (path) from the first activated astrocyte to the last one present in the subsection. Finally, the ensemble of co-activation paths are used to construct a graph which represents the functional connectivity of the underlying astrocytic network (Fig. 1D3), where each node of the graph is an astrocyte and an edge between two nodes (astrocytes) *A*_*i*_ and *A*_*j*_ carries a weight equal to the number of times they were consecutively activated in the recording section.

**Figure 1:**
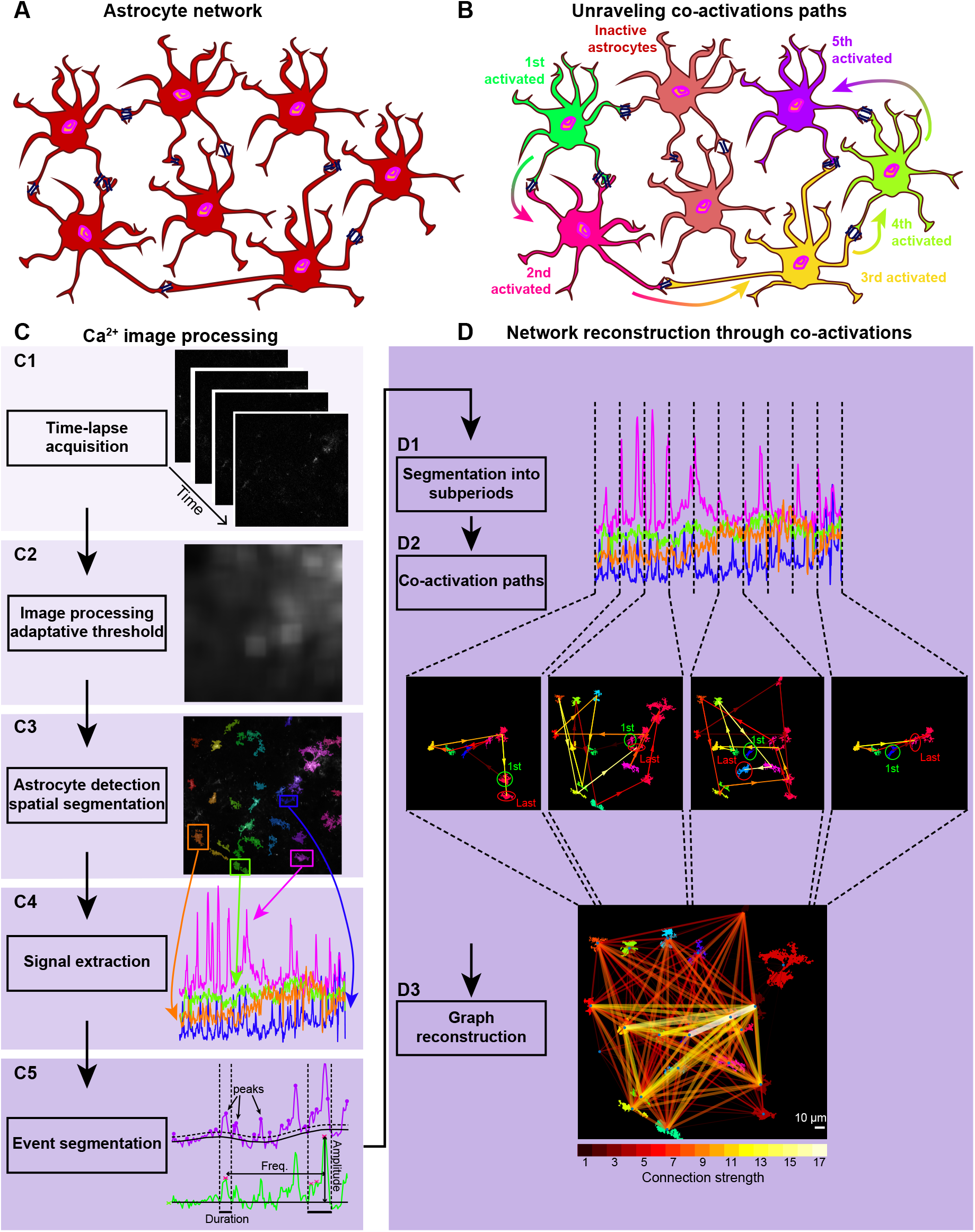
AstroNet pipeline. **A.** Schematic of an astrocyte network, connected through gap junctions. **B**. Schematic showing a hidden network path of sequentially activated astrocytes. **C**. Image processing steps: C1-movies are acquired from GCaMP6f Ca^2+^ signaling. C2-construction of an adaptive threshold for the segmentation step based on background luminosity. C3-Detecting all astrocytes (ROIs) with their characteristic shapes using a binarization on the sum of all images (over time). C4-Extracting the Ca^2+^ signals from astrocytic ROI. C5-Time-series segmentation into activation events and quiet inter-event periods (pink crosses highlight the peaks of the example events situated between the vertical dotted lines)4. **D**. Network reconstruction: D1-Ca^2+^ recording segmentation into subperiods (vertical dashed lines). D2-Activation path reconstruction for each subperiod. D3-Reconstruction of an astrocyte network from the recurrence direct co-activations of astrocytes.

### 1.1 From astrocytic burst detection to co-activation paths

Astrocytes exhibit spontaneous calcium activity [10, 20] (movie SI1) leading to the successive activation of various neighboring cells. In order to study whether these successive activations follow a repetitive order over realizations, we extracted from the calcium time-series the successive activated astrocytes that define a co-activation path. To do so, the first step is to extract and to segment the individual activity of each astrocyte (Fig. S1A). Due to the calcium baseline fluctuation (Fig. S1B-C), we first corrected these slow fluctuations (Fig. S1D-E and see Methods) to extract the individual astrocytes dynamics (Fig. S1F-G).

Once the individual dynamics have been identified, we collected the sequential activated astrocytes, to build the co-activation path. To this end, we segmented each recording session in *N* subperiods (Fig. 2A-C). Within each of these subperiods, activated astrocytes are ordered following the time of their main activation peak (Fig. 2D). This procedure allows to construct the activation path followed during this subperiod (Fig. 2E). Finally, we obtain *N* activation paths per recording session, that we used to reconstruct the graph of the astrocytic network (as described in the following section). Interestingly, the reconstruction procedure turns out to be relativity independent of the number *N* of subperiods chosen as shown in Fig. S3 (see sections 1.3 and Methods 3.7 for details).

**Figure 2:**
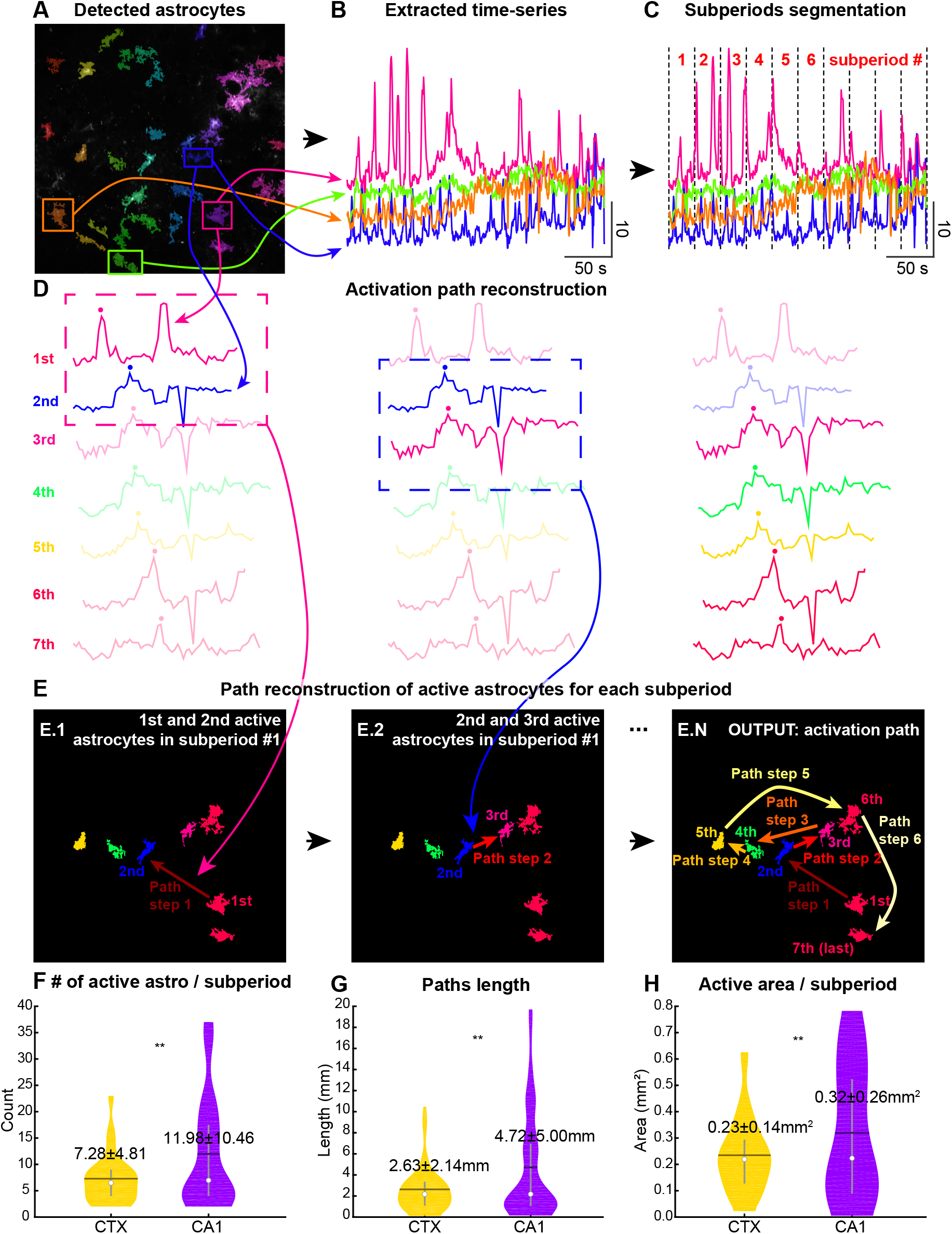
From astrocytic segmentation to co-activation paths detection. **A.** Detection of astrocytes ROIs (colored surfaces) obtained from single recordings. **B**. Time-series extracted from the detected astrocytes (color coded). **C**. Recording segmentation into *N* = 10 subperiods. **D**. Calcium time-series of individual ROIs (colored curves), ordered by their first peak time (colored dots) in one subperiod. **E**. Activation times (E.1-E.2) appearing in the field of view to reconstruct the activation path (red to yellow arrows). Output (E.N): one path is a collection of arrows indicating astrocyte activation order for one subperiod. **F-H**. Number of active astrocytes, paths length (sum of arrows lengths) and active area (surface of the convex hull of active astrocytes) for all subperiods in the CTX (yellow) and CA1 area of the hippocampus (purple).

When we applied this reconstruction procedure, although we detected a similar number of astrocytes in both regions (CA1 and CTX), we reported fewer activated astrocytes within subperiods in the CTX (7.28±4.81, Fig. 2F, yellow *n* = 8 slices) than in the hippocampus (11.98±10.46, purple, *n* = 9 slices, stars always indicate 2-sample Kolmogorov-Smirnov tests with ∗ 0.05 ≥ *p* ≥ 0.01, ∗∗ 0.01 *> p* ≥ 0.001, ∗∗∗ 0.001 *> p*). Similarly, the path length defined as the sum of the distances between consecutive astrocytes on the path is shortest in the cortex (2.63±2.14 mm, Fig. 2G) than in CA1 with 4.72±5.00 mm. Finally, we evaluated the active area, defined as the surface of the convex hull of all activated astrocytes within each subperiod. The area follows the same trend (CTX 0.23±0.14 mm^2^ and CA1 0.32±0.26 mm^2^, Fig. 2H). At this stage, we can conclude that the co-activation paths, that represent the spatio-temporal flow of information between connected astrocytes, already reveal clear differences in the local functional astrocytic network structure between the two regions.

### 1.2 From co-activation paths to graph reconstruction

After collecting all the co-activation paths in a given recording session, we counted the number of times *n*_*ij*_, a calcium activation in ROI *j* occurs just after the one in ROI *i* (Fig. 3A). This procedure allows to construct a connectivity matrix *C* = (*n*_*i,j*_)_*i,j*∈[1,*nROIs*]_ between all pairs of ROIs in the network (Fig. 3B and Fig. S4). We thus extracted one connectivity matrix (or graph) for each recording session. This matrix captures the local functional network organization from which we will be able to extract statistics such as the most active and/or most connected ROIs (see below).

**Figure 3:**
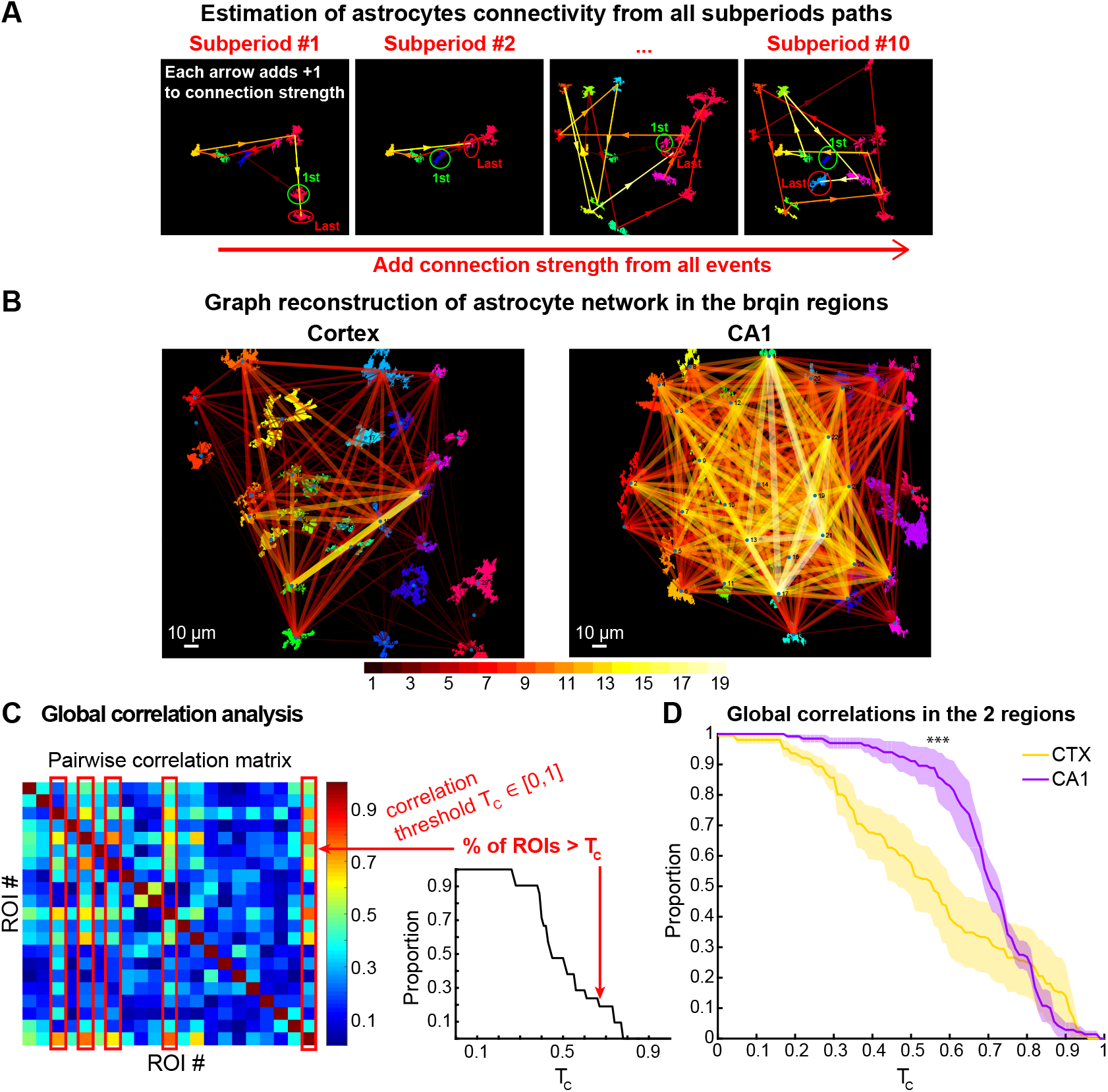
Astrocyte functional network reconstruction. **A.** Collecting activation paths from all the subperiods of a recording allows to evaluate the edges weights. **B**. Examples of reconstructed graphs, color coded according to the edges connectivity strength, showing two subgraphs with different connectivity levels: one much stronger (yellow edges) than the other (red edges). The size of the strongly connected yellow subgraph can vary from one region to the next (left, CTX vs right, CA1). **C**. Pairwise correlation matrix between all astrocytes time-series for one recording. **D**. Correlation analysis applied to the two regions: a faster decay reveals a less correlated network.

We then used this connectivity matrix to compare the level of local connectivity in the two regions CTX and CA1 area of the hippocampus, where one representative session per region is shown in Fig. 3B. A first estimation of the overall level of connectivity can be assessed from the pairwise Pearson correlation between the astrocytes’ signals over the entire session (Fig. 3C, left). To further quantify the correlation level between all ROIs, we constructed a correlation curve for each recording session by counting the proportion of ROIs that have at least one correlation higher than a specific threshold *T*_*C*_ that we vary between 0 and 1 (Fig. 3C, right). The closer to the top left corner is the curve, the more globally connected the astrocyte network is. We computed the area under the curve and obtained *AUC* = 0.56 ± 0.07 for the CTX and 0.70 ± 0.04 for the HPC thus revealing that the local astrocytic networks are more connected in the CA1 area of the hippocampus (Fig. 3D, purple), than in the CTX (yellow).

#### 1.2.1 Connection strength within the networks: number of direct neighbors

To further quantify the network connectivity, we counted the number of direct neighboring astrocytes of a given astrocyte/ROI, i.e. the number of other ROIs with which it has a direct connection. This number is the classical node degree (*deg*) in a graph (Fig. 4A, upper). We found that the CTX is characterized by a smallest mean degree (i.e. the smallest number of direct neighbors) with *deg* = 11.95 ± 5.52, number of slices *n* = 8), while the CA1 astrocytes are more connected with *deg* = 20.34 ± 13.33, *n* = 9). Interestingly, the distributions of node degrees (Fig. 4A, lower) confirm that trend: CA1 astrocytes have mostly a high connectivity level (45% of the ROIs have a degree ≥30, purple distribution), while 37.5% of the CTX ROIs have a connectivity degree lower than 11 (yellow). To conclude, these results support different levels of astrocytic network connectivities, higher in CA1 and lower in the motor cortex, as was previously reported for other regions (Hippocampus vs somatosensory cortex) based on connexin expression level [1].

**Figure 4:**
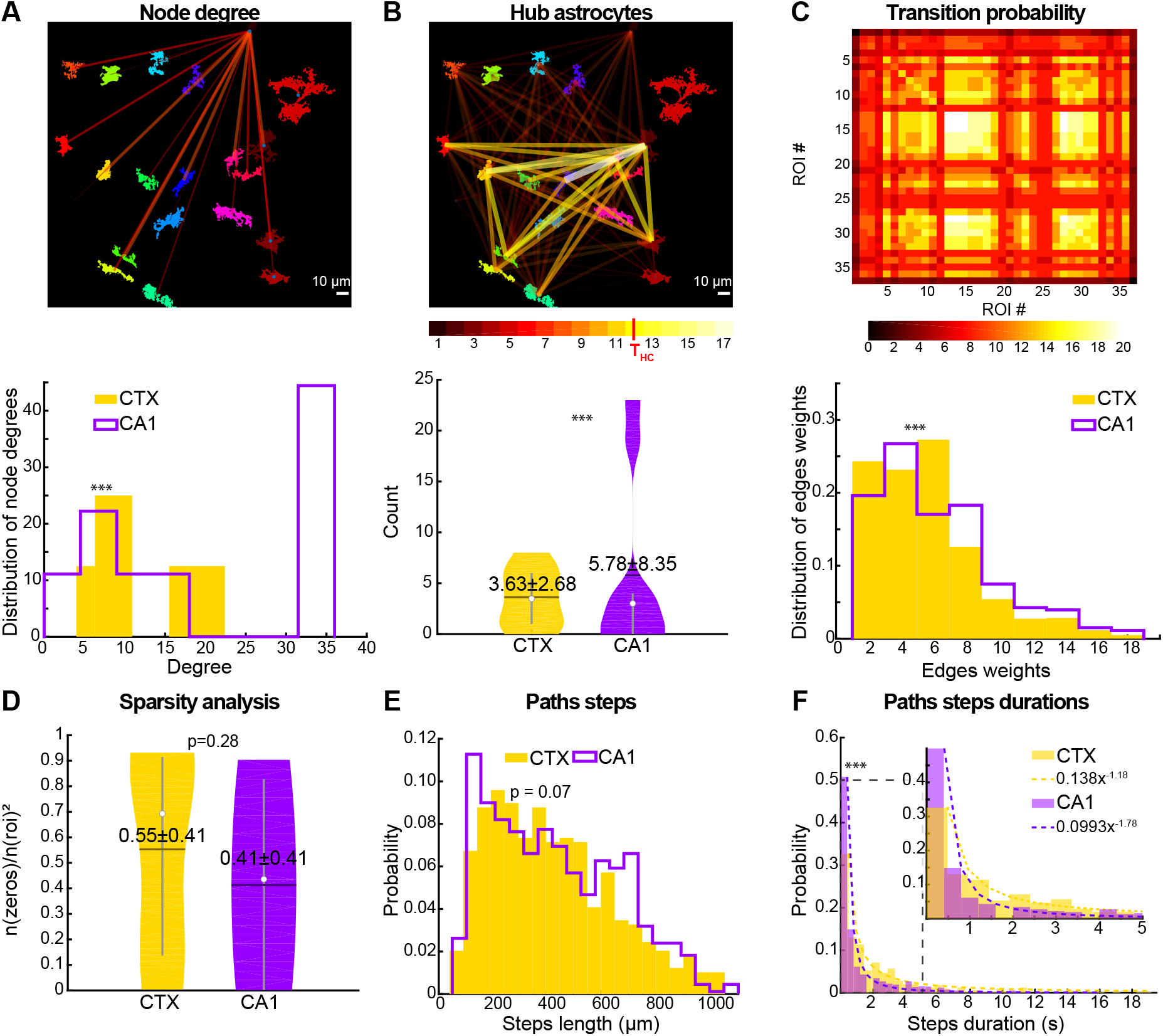
Statistics of the functional astrocytic network. **A.** Mean node degree (number of direct neighbors): schematic of the count of neighbors for one astrocyte (upper) and distribution (lower) in the CTX (yellow) and CA1 (purple). **B**. Number of hub astrocytes (highly connected: yellow upper graph) and violin plots for the two regions (lower). **C**. Transition probability matrix between nodes: connectivity matrix (upper, from the CA1), the color indicates the connection strength and distribution of edges weights between nodes (lower). **D**. Sparsity value (i.e. proportion of zeros in the connectivity matrix) for the two regions. **E**. Distribution of the length of paths steps in *μ*m. **F**. Distribution of path steps durations in seconds with power law fit (dotted lines), inset on short times (black dotted rectangle).

However, to further investigate how the functional astrocytic networks obtained by AstroNet differ from the quantification of physical connections made by diffusion through gap junctions, we use a pipette filled with biocytin. We found that the size of the astroglial networks in the hippocampus and in the cortex was not that different (Fig. S5), indicating that the calcium correlation network that we reconstruct differs from the sole passive diffusion. Possibly another mechanism is responsible for the difference in astrocyte network activity that we uncover with the AstroNet reconstruction.

#### 1.2.2 Hub astrocytes and highly connected sub-network components

To further evaluate the level of astrocytic connectivity, we decided in each graph, to focus on astrocytes with high connectivity level. We previously showed hetergeneous distributions, where some astrocytes were more connected than others (yellow subnetwork in Fig. 3B and Fig. 4B, upper). The nodes of these highly connected subgraphs can be considered as hub astrocytes, i.e. astrocytes that are participating in almost all activation events of the local network. We choose to count astrocytes that participate in at least 60% of events (Threshold *T*_*HC*_ = 0.6 ∗ 2*N* for undirected graph, where 2*N* is the maximal strength of an edge). We found that the CA1 area of the hippocampus has more hub astrocytes, with an average of 5.78 ± 8.35, or 17±24%, purple (Fig. 4B lower, purple)) compared to the cortex which is the less connected (3.63 ± 2.70 representing 28 ± 24%, yellow). We note that, although the absolute number of hub astrocytes is higher in CA1 compared to CTX, the proportions are reversed. This suggests that, in the less connected motor cortex, there is a larger proportion of astrocytes that need to participate in most activation events, however, in total the network remains sparser and less connected as we consistently saw with all the different features that we computed. Finally, the functional relevance of these hub astrocytes remain an open question, they could simply emerge as a consequence of extreme statistics of randomly connected networks [23] or could have a genetic origin, a question that should be further investigated.

#### 1.2.3 Transition probabilities between astrocytes

To further quantify the overall level of connectivity, we computed the distribution of the edges weights *n*_*ij*_ within each graph (Fig. 4C, lower). These weights can be understood as the transition probability from ROI *i* to *j*, i.e. the probability that if ROI *i* is activated, then *j* will be the next one. These values are also the components of matrix *C*, (Fig. 4C upper) and they differ significantly between the regions: again, the strongest weights are found in the CA1 area of the hippocampus (4.00 ± 4.19, purple), while the CTX has the lowest weights (1.57 ± 3.04, yellow), confirming once again the strongest connectivity in the CA1 hippocampus area.

#### 1.2.4 Astrocytic network sparsity

Conversely, we can also quantify the strength of the astrocytic network, by evaluating its sparsity, which is defined as the percentage of zeroes in the matrix *C* (Fig. 4C, upper). By definition,

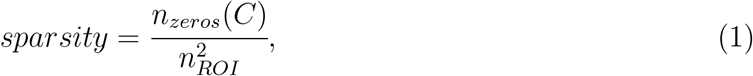

where *n*_*zeros*_(*C*) is the number of zeroes, normalized by the size of the network 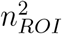, i.e. the number of ROIs that have been detected in a recording session. We found that CA1 hippocampal area has a lower sparsity (Fig. 4D 0.41 ± 0.41, purple) than the CTX (0.55 ± 0.41, yellow). However, in both cases we found two equally distributed subgroups, one characterized by sparse connectivity and the other one by dense connections. This disparity is reflected in the wide distributions of Fig. 4D and the lower significance of the statistical test results. At this stage, this intra-region difference remains unclear. To conclude, despite this variability, the sparsity analysis brings complementary information about the network compared to degree and hub analysis performed above. Furthermore, it confirms the trend seen all along that the functional connectivity is stronger in the CA1 hippocampus area than in the motor cortex.

#### 1.2.5 Statistics for two consecutively activated astrocytes

We computed the distribution of distances between each directly connected pair of astrocytes. This distribution is obtained from the length of the arrow connecting these two astrocytes in the co-activation paths. We found that the average distance is larger in CA1 (Fig.4E, purple, 430 ± 7.98 *μ*m) than in the CTX (418 ± 10.34 *μ*m). The distributions reveal two major differences between CTX (yellow) and CA1 (purple): there is a peak in CA1 for short paths steps (*<* 200*um*), which is absent in the case of the CTX, indicating a higher density of close neighbors with direct co-activations. A second peak in the CA1 distribution is situated at longer path steps (∼ 600 − 700*um*), also absent in the CTX indicating long-range direct connections. On the contrary, the distribution of CTX paths steps is more peaked at the medium lengths (∼ 200 − 500*um*), showing local variation of territories of astrocyte between CTX and CA1.

In parallel, we computed the duration of these consecutive steps (Fig. 4F), i.e. the time delay between the activation peak of astrocyte *i* and astrocyte *j* (the two extremities of the arrow). The distributions of these durations are well fitted by a power law of the form *y* = *Ax*^−*a*^, where *A* and *a* are parameters to be fitted. We found a significantly faster decay in the CA1 hippocampal area (*y* = 0.099*x*^−1.79^, purple) compared to the CTX (*y* = 0.14*x*^−1.18^, yellow). Finally, we summarize the main network quantification results in Table 1

**Table 1:** Summary of network connectivity parameters. (Figs. 2 & 4).

| Brain Regions | # Active nodes | # Hub nodes | Mean degree | Path length (mm) | Active area (mm<sup>2</sup>) |
| --- | --- | --- | --- | --- | --- |
| CTX | $7.28 \pm 4.81$ | $3.63 \pm 2.70$ | $11.95 \pm 5.52$ | $2.63 \pm 2.14$ | $0.23 \pm 0.14$ |
| CA1 | $11.98 \pm 10.46$ | $5.78 \pm 8.35$ | $20.34 \pm 13.33$ | $4.72 \pm 5.00$ | $0.32 \pm 0.26$ |

### 1.3 Stability analysis of network reconstruction

#### 1.3.1 Convergence of the network parameters

To ensure that the graph reconstruction and analysis of the astrocytic network from the calcium time series is stable, we developed a convergence analysis, which consists in computing various network properties while varying the number of subperiods *N* ∈ ⟦1, 20⟧. These network properties are: the mean and variance of the graph’s node degree, the mean path length, the number of hub astrocytes, the active area per subsection and the number of active ROI per subsection (Fig. S3). Interestingly, after we ran the AstroNet pipeline analysis for increasing *N* (Methods section 3.7 and Fig. S3), we obtained a convergence towards the final value already after *N* = 5 in most cases. Indeed, for *N* too small, there is not enough paths to reconstruct a robust graph, as soon as *N* is big enough, the networks properties do not depend on the value of *N* (Fig. S3). At this stage, we concluded that with *N* = 10, the AstroNet algorithm provided a robust reconstruction of the underlying astrocytic network.

#### 1.3.2 Stability of the astrocytic network revealed by a time-lapse analysis

To investigate the stability of the astrocyte networks obtained with AstroNet over a period of few minutes, we conducted a time-lapse analysis. We reconstructed and compared the network reconstructions at two different times separated by 5 minutes (see methods section 3.7): we recorded successively at the same position in the same slice (Fig. 5A, timepoint 1, upper and timepoint 2, lower) for 5 min intervals, and then built and compared the reconstructed graphs of the local network for the two timepoints (Fig. 5B) with their corresponding connectivity matrices (Fig.5C). Fig. 5A-D shows the results for one example in CA1, we show other graphs both in the CTX (yellow, n=2) and CA1 (purple, n=6) in Fig. S6.

**Figure 5:**
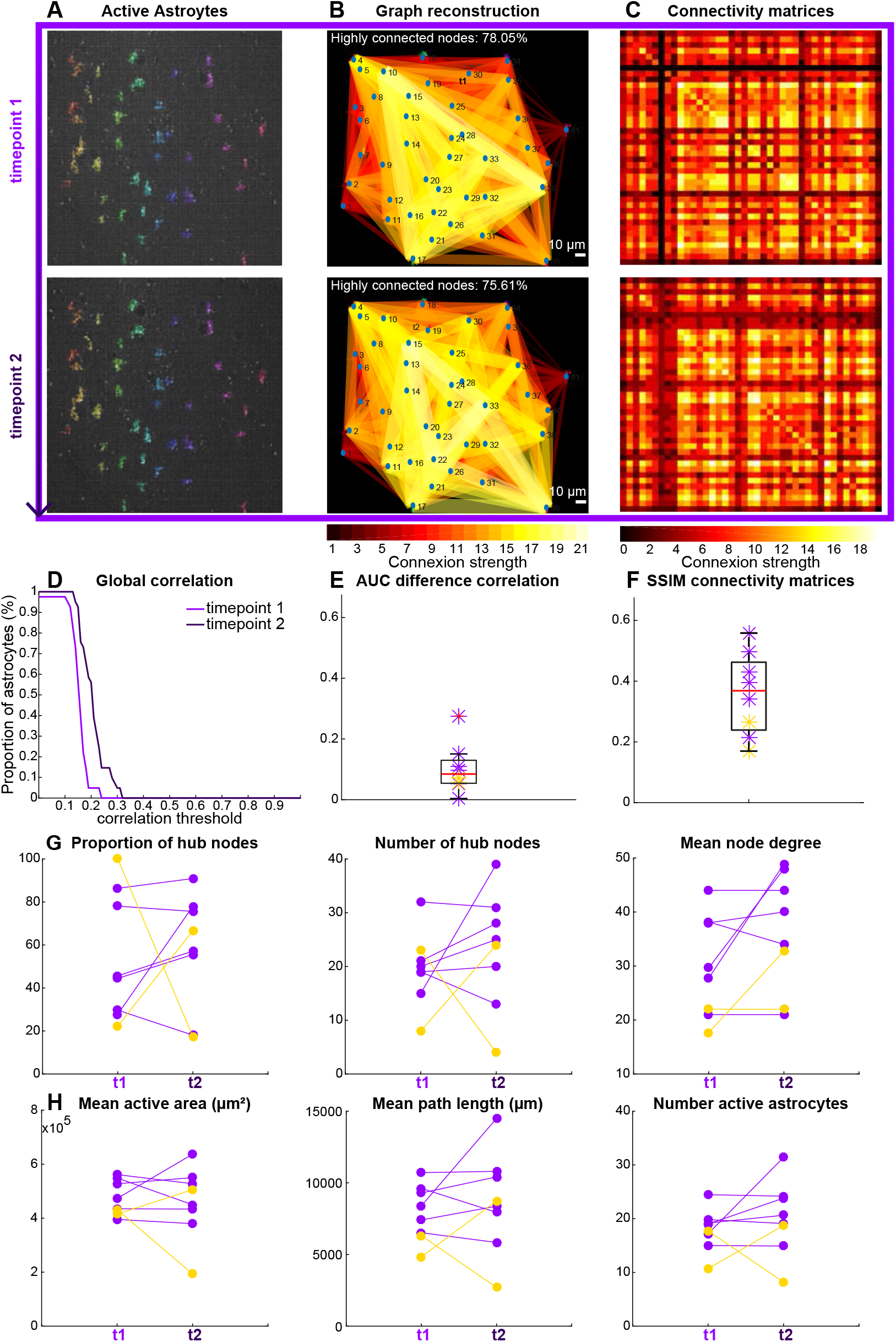
Stability of AstroNet pipeline based on time-lapse experiments. **A.** Recording at timepoint t1 (upper) and timepoint t2 (lower) overlayed with the detected ROIs recorded from calcium imaging in a CA1 slice. **B**. Reconstructed graphs at time t1 (upper) and t2 (lower). **C**. Corresponding connectivity matrices. **D**. Correlation curves at t1 (light purple) and t2 (deep purple). **E**. Difference between the AUC at t1 and t2 (absolute value) for the n=2 CTX slices (yellow) and n=6 CA1 slices (purple). **F**. SSIM13between the connectivity matrices at t1 and t2. **G**. Graph features at t1 (left points) vs t2 (right points): from left to right: Proportion of hub ROIs, number of hub ROIs, Mean node degree. **H**. Activation path features at t1 vs t2: from left to right, Mean active area (*μm*^2^), Mean path length (*μm*) and number of active astrocytes.

To quantify the variations in the global correlation between the two timepoints t1 and t2, we compared the correlation curves between the two timepoints (Fig. 5D) by computing the differences between the AUC of both curves for all pairs of recordings. We found that, for all pairs, the difference was small (Fig. 5E, 0.10 ± 0.08). Furthermore, to quantify the differences in the reconstructed graphs, we computed the Structural Similarity Index Measure (SSIM) between the connectivity matrices of timepoints t1 and t2 for all pairs of time-lapse recordings (Fig. 5F) and obtained 0.36 ± 0.13. We found that in most recordings the SSIM index is higher than 0.4, indicating a strong similarity [24]. Finally, we extracted the same statistical properties of paths and network that we computed in the previous section, revealing the proportion and number of hub astrocytes, mean node degree, mean active area, mean path length and number of active astrocytes for each pair of recordings (Fig. 5G-H): we found that in most cases it remains stable from timepoint 1 to timepoint 2 specifically in the CA1 region. In CTX recordings, we report more variability in the mentioned parameters. This variability could originate from the different level of activity between the two timepoints that we observed.

The present results show that the reconstructed graphs are stable over time and that AstroNet is able to account for this property.

## 2 Discussion

We presented here a method for the reconstruction of local astrocytic functional networks from recordings of spontaneous calcium activity. This pipeline relies on detecting astrocytic bursting times within the field of view (size 100 *μ*m by 100 *μ*m), where calcium fluorescence signals were recorded, and ordering them as they are sequentially activated (see Fig. 2). The outcome of our approach is a collection of paths between astrocytes and a graph representing the local astrocytic functional network, encompassing 20 to 40 connected astrocytes. Interestingly, although astrocytes located at the periphery of the field of view may have connections outside of it and we may thus underestimate their connectivity level, most astrocytes are well within the image and therefore counterbalance this possible underestimation when it comes to extracting statistics for the whole astrocyte population. Thus, this sampling group of astrocytes is sufficient to reveal several statistics showing for example that there are fewer active astrocytes in the CTX compared to CA1 (7.28 ± 4.81 vs 11.98 ± 10.46 respectively).

The presented method and algorithms allowed us to extract various statistical properties of the astrocytic functional network, such as the degree of connectivity (number of direct neighbors), identification of the strongest connections across the network, identification of hub astrocytes, and the derivation of transition probability matrices to quantify path statistics.

The present algorithm complements softwares such as AQuA [21] that quantifies the ROI of calcium and neurotransmitter activity in fluorescence imaging datasets but does not reconstruct the underlying network. The new preliminary BioRxiv version called AQuA2, posted during the present submission, does allow for ROI segmentation, similar to AstroNet (see SI for a comparison), but does not integrate the astrocyte network reconstruction and connectivity quantification, which is the main focus of AstroNet.

Many software tools exist for the analysis calcium activity in both glial and neuronal cells, including GECI-quant [25], Suite2P [26], LC Pro [27], CaSCaDE [28], CaImAn [29], and STARDUST [30]. While LC Pro and STARDUST are more glia-focused, making them better suited for analyzing astrocyte-specific calcium signals, other tools like Suite2P and CaImAn are primarily designed for neuronal calcium signals and are not inherently tailored for astrocyte connectivity analysis. However, unlike the approach introduced in AstroNet, none of these tools provide direct estimation of astrocyte functional connectivity. AstroNet is specifically optimized for detecting and quantifying astrocyte calcium signal correlations, including spatio-temporal relationships that reflect network connectivity. In a different direction, identifying sub-graphs with stronger connectivity enabled differentiation between types of networks, from highly connected small worlds [31] to more uniformly connected structures, as measured by the sparsity of local connections (Fig. 4).

Using this computational pipeline, we compared astrocytic connections within two different brain regions: the motor cortex (CTX), and the hippocampus (CA1). We reported differences in the connectivities of the local astrocytic networks, with the hippocampus displaying the highest density of connections (mean degree of connectivity ≈ 20) and the cortex the lowest (≈ 12) (see Table 1). The higher variability in CA1 compared to CTX indicate that the different parameters we extracted (Table 1) are more preserved across slices in the CTX. Similarly, we reported here larger distributions in CA1 for the extracted statistics (Fig. 4A-C and E) compared to the CTX. These differences were also reflected in the global correlation levels between astrocyte time-series (Fig. 3D) and the number of highly connected astrocytes (hub astrocytes), averaging 3.6 in the cortex vs 5.8 in CA1 (Fig. 3B, yellow structures, and Fig. 4B, left). While the role of GABAergic hub neurons in orchestrating synchrony in developing hippocampal networks has been established [32], the function of hub astrocytes remains unclear.

In conclusion, distinct brain regions appear to have specific astrocytic connectivities characterized by their degree of connectivity, overall connectivity, or the presence of highly connected sub-regions. These results may reflect the reported heterogeneity in gap junction channel abundance (connexin expression) and/or permeability, indicative of anatomical connectivity [33]. It could also reflect differences in astrocyte number and morphology, as the hippocampus, with its diverse astrocyte morphologies, contains a high number of astrocytes compared, for example, to the somatosensory cortex, where astrocytes are more uniformly distributed. Additionally, we found that the mean path length (total path distance between the first and the last activated astrocytes, Fig. 2) was higher in the CA1 hippocampal area, averaging at 4.7mm compared to only 2.6mm in the cortex (Table 1).

The present approach allows us to quantify the local level of connectivity from spontaneous calcium transients in astrocytes, addressing the growing need to decode astrocyte signaling and its relevance to brain function [34]. While the method does not detect strictly repeating activation paths (no deterministic causal sequences), a high correlation was observed between specific subgroups of neighboring astrocytes that were more connected to each other than the others, despite their similar distances (Fig. 3B, yellow subgraphs, and Fig. 4B). This result suggests a direct mechanism of activation between these specific astrocytes. Dendritic or neuronal activity is much faster than astrocytic calcium dynamics, thus it is unlikely that the long paths we found here that are usually recurrent (coming back to the same astrocytes) and lasting up to a few seconds could be the results of co-activated neurons. Relying solely on passive diffusion through gap junctions would predict a uniform co-activation pattern, which our analysis disproves. The local enriched connectivity reported here raises questions about how this information can clarify neuron–glia interactions in specific networks [35]. The present analysis is based on soma calcium events and disregard protrusion-based activity, which would require a more refined analysis.

Finally, it is worth noting that this pipeline could also be applied to analyze neuronal fluorescent imaging. Parameters related to time and space scales can be adjusted to fit cell signals at various time-scales. The modularity of the implemented method also allows for the use of any sub-section of the signal. For instance, it is possible to use calcium activation times as an input for the graph reconstruction method, yielding graphs of local neuronal connectivity for a relatively slow signal compared to spiking activity. Differences in the structure of these graphs could provide quantification of local changes occurring at an intermediate scale between single-cell analysis and wider brain areas. Additionally, the present approach could be generalized to in vivo one-photon imaging, allowing the tracking of ROIs in freely behaving animals. The most challenging step in this case would be ROI detection due to the coarse resolution of one-photon imaging.

## 3 Methods

The method section is organized as follows: we first recall the experimental settings used to collect calcium fluorescence recordings in the two brain regions. Second, we describe the image processing steps used to detect the astrocytic ROIs and extract segmented time-series. Third, we introduce the signal processing steps to extract the astrocytes activation events. Then, we introduce our algorithm to reconstruct astrocytic networks based on consecutive calcium burst co-activation sequences. Finally, we describe the stability analysis and corresponding time-lapse experiments.

### 3.1 Experimental protocol

#### 3.1.1 Ethics on Animal manipulation

All experiments were performed according to European Community Council Directives of 01/01/2013 (2010/63/EU) and of the French ethic committee (ethics approval delivered by the French ministry of higher education, research and innovation). Experiments were carried out using the GFAP-CreERT2-GCaMP6f transgenic mouse line expressing the GCaMP6f calcium indicator in astrocytes. This mouse line was obtained by crossing the GFAP-creERT2 line with the Cre-dependent B6J.Cg-Gt(ROSA)26Sortm95.1(CAG-GCaMP6f)Hze/MwarJ transgenic mouse line (referred as Ai95(RCL-GCaMP6f)-D (C57BL/6J)), expressing a floxed-STOP cassette blocking transcription of the fast calcium indicator GCaMP6f (The Jackson Laboratory, Stock No: 028865), and which were previously characterized [36]. Tamoxifen (10 mg/ml in corn oil, Sigma) was injected intra-peritoneally (100 mg/kg body weight) during four consecutive days at around postnatal day 30 and experiments were performed at least three weeks after the last injection. Mice of both genders were used at postnatal days 50 to 60, and were housed under standard conditions with ad libitum access to food and water.

#### 3.1.2 Two-Photon calcium imaging

Acute transverse hippocampal slices (300-400 *μ*m) were prepared as previously described [37] from 50-60 days-old mice from WT (C57BL6) mice. GCaMP6f mice aged 50-60 days were decapitated, and their brains were rapidly extracted and submerged in cold (0-4^°^C) slicing artificial cerebrospinal fluid (aCSF) containing (in mM): 119 NaCl, 2.5 KCl; 2.5 CaCl2, 26.2 NaHCO3, 1 NaH2PO4, 1.3 MgSO4, 11 D-glucose (pH = 7.35). This solution was saturated with a gas mixture of 95% O2 and 5% CO2. Coronal brain slices, 400 *μm* thick, were cut using a vibratome (VT1200S; Leica) targeting slices around Bregma 0 mm for cortical recordings, and around Bregma −1.7 mm for hippocampal recordings. Slices were then transferred to a continuously oxygenated (95% O2–5% CO2) holding chamber containing aCSF for at least 1h before recording.

Slices were transferred to the recording chamber mounted on an Axio Examiner Z1 microscope (Carl Zeiss) equipped with a two-photon system (3i, Intelligent Imaging and Chameleon laser, Coherent) and were perfused with ACSF at a rate of 4 ml/min. Experiments were performed in the presence of picrotoxin (100 *μ*M) and a cut was made between CA1 and CA3 to prevent recurrent electrical activity between the two regions. Astrocytes expressing AAV GCaMP6f were excited at 920 nm and fluorescence detected after a 525/40 emission filter and a 580 nm dichroic mirror. Water-immersion objectives of either 20x, NA 1.0 or 40x NA 1.0 (Carl Zeiss) were used. Images were acquired at 3 Hz through a 40^°^ water immersion objective (NA 0.95, Olympus). Images were pre-processed off-line with Slidebook imaging software (3i) by applying a median filter.

##### Astroglial gap-junction coupling

To visualize astroglial networks, biocytin (7 mg/ml, Sigma) was included in the intracellular solution, allowing it to diffuse throughout the network over 20 minutes. The slices were then fixed overnight in 4% paraformaldehyde (PFA), followed by incubation in phosphate-buffered saline (PBS) containing 1% gelatin and 1% Triton X-100 (PGT 1%). Visualization was achieved using Alexa Fluor 555-conjugated streptavidin (1:300 in PGT1%, Invitrogen). After several PBS washes, the slices were mounted in Fluoromount-G (Southern Biotechnology) and examined using a confocal laser-scanning microscope (Leica DMI6000 Inverted SP5) equipped with a 20×/0.75NA objective. Z-stacks of consecutive confocal images were captured at 0.5 *μm* intervals using a 561 nm Diode-pumped solid-state (DPSS) laser controlled by LAS AF software (Leica). Cell counting was performed with ImageJ software.

##### Data acquisition

We acquired data under standardized conditions with a scanning rate of 3 Hz, at a depth of 50 microns below the surface, with consistent laser power. Changing the laser power could destabilize astrocytes excitation. This standardization is essential for ensuring the reproducibility of the present datasets. We used 5 min recordings and performed the analysis on the entire recordings that we further split into 10 subperiods for the network reconstruction analysis (section 3.4). Our recordings and codes are available on Zenodo, github and on the lab’s website www.BioNewMetrics.org, see below.

### 3.2 Principle of spontaneous calcium activity in astrocytes

Calcium spontaneous activity in astrocytes is generated by calcium-induced-calcium-release (CICR) from the endoplasmic reticulum (ER) triggered by the opening of Ryanodin receptors (RyRs) and/or IP3-receptors [20, 38, 2]. The up-stream signaling can be generated by neuronal activity or random calcium events with IP3 transients in astrocytes. Once calcium ions are present in the cytoplasm, they can either be pumped back into the ER or diffuse to the neighboring astrocytes through gap junctions. These calcium transients generate a bumped shape in fluorescent time-series 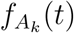, recorded in a region of interest (ROI) around astrocyte *A*_*k*_. We build the present pipeline based on the hypothesis that calcium transients can be detected at a time *t*_*k*_ in a given epoch by detecting the global maximum (within this epoch) of the calcium fluorescence intensity for a time series. In an epoch going from 0 to time T, this is given by:

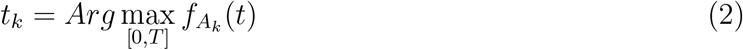

Network events consist of ordered time sequences (min(*t*_1_, ..*t*_*m*_), .., max(*t*_1_, ..*t*_*m*_)) among detected astrocytes. The difference between two consecutive time points 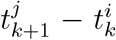 for two astrocytes *i, j* presenting a consecutive calcium burst, reflects calcium signal propagating from astrocyte *i* to *j*.

### 3.3 Astrocyte detection and events segmentation

#### 3.3.1 ROI detection

The ROI detection requires two steps:

##### 1. Signal summation and normalization

To detect the position of all active astrocytes in each recorded session, we sum all frames over time, resulting in a combined image *I*_*sum*_, that we normalize as follows:

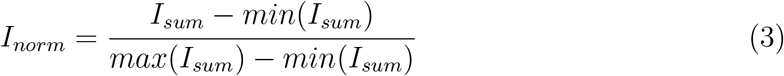

Thus pixel values *I*_*norm*_ are now in the [0, 1] range.

##### 2. Image binarization

To account for the heterogeneity of the luminous intensity in the image (due to the laser’s position), we use an adaptive threshold *T*_*adapt*_ obtained with the built-in MATLAB function *adaptthresh* with a threshold parameter *T* = 0.5. We then binarize the normalized summed image *I*_*norm*_ with this threshold *T*_*adapt*_. Finally, we remove all detected ROIs that are too small to be an astrocyte soma (smaller than a size *T*_*min*_ = 30 pixels). This allows us to filter artifactual detections due to the presence of noise in the recordings. All the remaining regions are kept as potential ROIs. The threshold values *T* and *T*_*min*_ are input parameters of the method and can be adjusted for each type of recordings.

#### 3.3.2 Calcium signal extraction and baseline correction

From each detected astrocyte *a*, we extract the temporal fluorescent signal by averaging the intensity over all pixels in the ROI (SI Fig. S1A) at each time frame *t* from the normalized data. To treat high resolution imaging, we can use a spatial down-sampling parameter, as explained in the SI. We obtain an activation time-series denoted *s*^*a*^(*t*) (Fig. S1B) for each astrocyte ROI *a*. We kept active astrocytes for which the calcium dynamics has a variation greater than 10% over time, compared to their mean value ⟨*s*^*a⟩*^.

We segmented the time-series to detect the calcium activation events and extracted their characteristics by correcting slow fluctuations of the baseline (SI Fig. S1C-E), that can vary from one ROI to the next, as well as the noise intensity of each signal. This correction is made in several steps:

##### 1. Peak detection

Calcium peaks in the fluorescent signal are detected using the built-in MATLAB function *findpeaks* with a minimal amplitude *minDepth* = 0.1 (Fig. S1C, purple dots).

##### 2. Baseline signal outside of peaks

We recover the baseline fluctuations of the signal, from the time intervals during which astrocyte *a* is inactive (Fig. S1D, red curve). To do so, we remove the duration Δ*t*_*ev*_ = 10s before and 1.5Δ*t*_*ev*_ = 15s after each detected peak leading to an incomplete output signal *r*^*a*^(*t*), only defined outside of the activation periods, i.e. *r*^*a*^(*t*_*inact*_ = *s*^*a*^(*t*_*inact*_)), where the timepoints *t*_*inact*_ are left after removing the activation periods. The function *r*^*a*^(*t*) is not defined during the activation periods.

##### 3. Complete baseline reconstruction

To construct a global baseline signal *b*^*a*^(*t*) for each astrocyte *a* over the entire time series, we fitted the signal *r*^*a*^(*t*) obtained from the previous step, with a smoothing spline (with smoothing parameter *p*_*smooth*_ = 7.10 − 5) using the MATLAB fitting tool (Fig. S1E, black curve). This procedure allows to define the baseline *b*^*a*^(*t*) for all times.

##### 4. Corrected calcium signal

The corrected signal (SI Fig. S1F, dark green curve) is obtained by subtracting the baseline *b*^*a*^ from the activation time-series *s*^*a*^, leading to 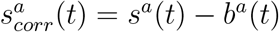.

The resting value of signal 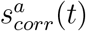 is now zero (Fig. S1F, solid black line). This signal is used to segment and extract calcium event statistics, as described in the next paragraph.

#### 3.3.3 Signal segmentation and statistics extraction

We segment the corrected signal 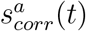 as follows:

##### 1. Event threshold definition

An event is detected if 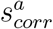 reaches a threshold *T*_*e*_ (SI Fig. S1F dotted black line) defined as *T*_*e*_ = *min*(4.5*σ*^*a*^, *T*_*high*_) where *σ*^*a*^ is the empirical standard deviation of the signal outside of peaks *r*^*a*^(*t*) and *T*_*high*_ = 0.08 is an fixed value chosen empirically to avoid too high event thresholds. This can happen when the signal outside of peaks *r*^*a*^(*t*), is defined on very few timepoints (i.e. for a very active astrocyte) and thus its empirical standard deviation would not be very representative of the actual noise of its signal. In all other cases the empirical variance *σ*^*a*^ is a good estimate of the noise amplitude of the signal.

##### 2. Event detection

An event is detected at time *t*_*d*_ if 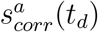 reaches the threshold *T*_*e*_ that we defined in the previous point. Then, the beginning of the event is set as the last time *t*_*i*_ *< t*_*d*_ where 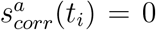 (i.e. at the baseline). The event end is defined as the next time *t*_*e*_ *> t*_*d*_ when the signal is back at its baseline value i.e. 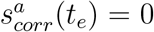 with *t*_*e*_ *> t*_*d*_.

##### 3. Events statistics extraction

The signal is then segmented into events (SI Fig. S1G, blue) and inter-events (dark orange) from which we extract the durations *d* (blue), the amplitudes *α* (black), the inter-event durations *i*_*e*_ (dark orange), the event frequencies *f* (red) and also the number of peaks *n*_*p*_ per event (light orange). This is done for every ROI.

### 3.4 Reconstruction of the local astrocyte network

We detail here the procedure that allows us to construct the connectivity graph between the detected astrocytes from the consecutive sequences of calcium events. This network reconstruction is based on the segmentation of the recordings in *N* subperiods over the whole time of the recording used to identify the calcium co-activation paths between astrocytes for each subperiod. This collection of *N* co-activation paths is then concatenated to build the global connectivity graph with estimated weighted edges for the whole recording session. We detail theses steps in the following sections.

#### 3.4.1 Detection of the co-activation paths

The recording sessions are first segmented into *N* subperiods of equal duration (Fig. 2C). We then study the co-activation dynamics within each subperiod as follows:

##### 1. Detection of the astrocytes activated during the subperiod *n* ∈ ⟦1, *N*]

An astrocyte *a* is considered active during the subperiod *n* if there is at least one time *t* in the subperiod *n* for which 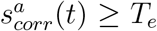, where *T*_*e*_ is the event threshold defined in the previous section.

##### 2. Estimation of the activation order during the subperiod *n*

We order the activated astrocytes with respect to the time of their main activation peak during the subperiod *n* (Fig. 2D, the colored dots indicate the main peak for each astrocyte’s signal within the subperiod).

##### 3. Co-activation path reconstruction

The co-activation path is obtained iteratively by joining the ordered astrocytes by arrows pointing toward the next activated astrocyte, going from the first activated to the last one (2E, red to yellow arrows). A such path is obtained for each subperiod, i.e. for one recording session we obtain *N* activation paths.

#### 3.4.2 Constructing a connectivity graph

A graph is an ensemble of *n*_*a*_ nodes connected by weighted edges. The present graph construction procedure is based on the concatenation of the co-activation paths described in the previous sub-section. Specifically, each astrocyte ROI *a* constitutes a node of the graph. The edge weights are then extracted from the co-activation paths in two ways leading to a directed or an undirected graph:

##### 1. Directed graph

built by adding the value +1 to the edge weight *n*_*i,j*_ of the connectivity matrix *C*, each time there is a direct arrow from astrocyte *i* to astrocyte *j* in the collection of co-activation paths obtained above. This leads to an asymmetric matrix of size (*n*_*ROIs*_*×n*_*ROIs*_) (where *n*_*ROIs*_ is the total number of active ROIs in the recording session). Each value is between 0 (no connection between this pair) and *N* (strongest connection). The result is an asymmetric connectivity matrix of the graph.

##### 2. Undirected graph

the procedure is similar, except that we add +1 to both *n*_*i,j*_ and *n*_*j,i*_ regardless of the direction of the arrow each time astrocytes *i* and *j* are directly connected by an arrow in a path. This results in a symmetric matrix of size (*n*_*ROIs*_ × *n*_*ROIs*_), where each value is between 0 and 2*N* (Fig. 3B).

To conclude, we obtain a connectivity graph *G* = (*A, C*), where *A* is the ensemble of vertices (i.e the atrocytes ROIs) and *C* is the connectivity matrix. In summary:

1. The ensemble of nodes associated to detected astrocytes are 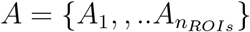.
2. The connectivity matrix 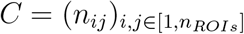 counts the number of times edge *i* → *j* is activated in an event.

We define the probability density matrix from the connectivity matrix by dividing each coefficient by the sum of the coefficient in a given raw.

### 3.5 Pearson correlation analysis

The global correlation analysis (Fig. 3C-D) is performed using the Matlab built in function *corr*. This function computes the pairwise Pearson correlation coefficient between each pair of signals (i.e. for each ROI). The Pearson correlation coefficient *ρ* between two time-series *X* and *Y* is defined as

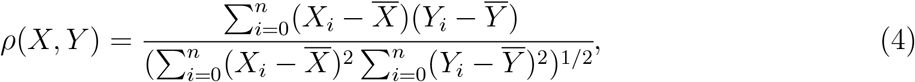

where 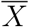 denotes the average value of *X*.

### 3.6 Highly connected subgraph and hub astrocytes detection

We can define a subgraph (*G*_*max*_, *C*) that contains only the most connected nodes (see Fig. 3B, yellow subgraph). This is the subgraph where all edge weights *n*_*ij*_ are higher than the threshold *T*_*HC*_. Here we show the statistics for *T*_*HC*_ = 0.6 max(*C*) i.e *n*_*ij*_ = 12 in the case of the non-oriented graph (see Fig. 3B, yellow subgraphs, and Fig. 4B). Subgraphs of any connectivity level could be extracted by varying the value of *T*_*HC*_ which is an input parameter in the function *activGraph* that constructs the graphs and gets its statistics (SI, section 1.5).

The hub astrocytes are the nodes of the highly connected subgraph, i.e. the ones connected to at least 60% of the network (i.e. of degree *d* ≥ *T*_*HC*_).

### 3.7 Stability analysis

#### 3.7.1 Effect of the number of subsections on the graph reconstruction

To guarantee the stability of the network construction algorithm, we studied how the choice of the number of subsections *N* affects the graph’s properties. To do so, we varied the number *N* of subsections and ran the whole pipeline. Specifically, we ran the entire AstroNet pipeline for all the available recordings from the two brain regions with *N* ∈ ⟦0, 20⟧. We then plotted several extracted features that quantify the network organization vs the value of *N*. These features are: mean node degree, variance of node degree, mean path length, number of hub astrocytes, active area per subperiod and number of active astrocytes per subperiod. We obtained one curve per recording, for each observed parameter vs the values of *N* (Fig. S3).

#### 3.7.2 Time-lapse analysis for network reconstruction stability

To test the stability over time of AstroNet, we recorded in the same region, two successive recordings of 5 min, with a resting period of also 5 min, while the slice incubated in ACSF oxygenated with later closed.

To detect all active ROIs, we concatenated the two recordings and ran the ROI detection step. To quantify the changes between the two recordings, we computed the difference between the area under the curves (AUC) of the global correlation curves of both recordings. To compare the network structure, we used and computed the Structural Similarity Index Measure (SSIM) [39] between the two connectivity matrices (Fig. 5C, F). The SSIM offers a trade-off between direct difference measurements such as Euclidian distance or Mean Squared Error (MSE), and indirect ones such as correlation [24]. A SSIM of 1 means total similarity and 0 total difference. Using this measure for matrix comparison, an SSIM greater than 0.4 is considered as a strong similarity [24].

### 3.8 AstroNet MATLAB toolbox

AstroNet is available at https://zenodo.org/records/13889962 and https://github.com/louzonca/NetConstruct and www.bionewmetrics.org. The pipeline (described in 1), is modular made of independent functions for each of the steps. This allows to process only part of the pipeline at a time. For example, one could provide time-series of any type of cellular activation (e.g. neuronal patch clamp or MEA recordings or calcium time series of various types of cells) and start AstroNet at the segmentation step (Fig. 1C5) to extract the local functional network properties. The contents, scripts and functions are described in the SI (section 1).

## Acknowledgments

LZ was supported by Fondation pour la Recherche Medicale (FRM FDT202012010690). DH is supported by ANR AstroXcite, Memolife and the European Research Council (ERC) under the European Union’s Horizon 2020 research and innovation programme (No 882673). NR is supported by ANR (AstroXcite) and Memolife.

