## Supplementary information for "Unveiling the Functional Connectivity of Astrocytic Networks with AstroNet, a Graph Reconstruction Algorithm Coupled to Image Processing"

We detail here how to use the AstroNet Matlab toolbox and the various functions used for the signal and image processing steps. We then describe some supplementary results about the statistics of the individual bursting events and the stability analysis for the graph reconstruction with respect to the number of segmented subsections chosen $N$. We the display six supplementary figures. Finally we provide a detailed comparison of AstroNet vs AQuA2 and other existing calcium analysis tools.

### 1 AstroNetConstruct toolbox description

The AstroNetConstruct toolbox contains various functions performing the unitary steps of the entire analysis pipeline (Main Fig.1). We detail here each function. The toolbox also contains a script example that runs the entire analysis at once.

#### 1.1 Image processing and ROI detection

In the toolbox, the image processing and ROI detection step is performed by the function *cellDetectFun* that is used as follows:

```
[cellMap,nROI] = cellDetectFun(data,T_min,T);
```

where, **data** is the raw calcium recording (that can be provided in any MATLAB supported video format), $\mathbf{T}_{min}$ is the minimal size in pixels for a detected ROI, and **T** is the adaptive binarization threshold parameter. The function returns the outputs: **cellMap** an image the size of the data frames giving the position of the detected ROIs and associating them with a label number, and **nROI** the total number of ROIs detected.

[1]Applied Mathematics and Computational Biology, IBENS, Ecole Normale Supérieure, PSL University, Paris, France.
[#] Present address: Center for Brain and Cognition, University Pompeu Fabra, Barcelona, Spain.
[2]Neuroglial Interactions in Cerebral Physiology and Pathologies, Center for Interdisciplinary Research in Biology, Collège de France, CNR UMR 7241, INSERM U1050,PSL, Paris, France.
[3]Churchill College, University of Cambridge , CB30DS UK.
Co-corresponding

#### 1.2 Calcium signal extraction and baseline correction

The calcium signal extraction is implemented using the function *extractIndivSignals* as follows:

```
[decharges,dechNorm,varList] = ...
extractIndivSignals(data,cellMap,dSiz,ROI_list,varMin);
```

where, **data** is again the raw calcium recording, **cellMap** is the map of detected cells obtained as the output of *cellDetectFun*, **dSiz** is a spatial down sampling factor (a positive integer) introduced for efficiency ($dSiz = 1$ means no down sampling and the higher the value the more down sampling), $\mathbf{ROI}_{list}$ is the list of the ROI labels that we would like to extract the signal from (we use all of them in the results of the main text), **varMin** is the minimal variation level – in percents of an ROI signal compared to its mean value – needed to consider the ROI as active (here we use $varMin = 10\%$). The outputs are: **decharges** a matrix of the size of the total duration (in frames) of the recordings times the number of considered ROIs that contains the extracted time-series for each ROI, *dechNorm* which is the same as decharges but with signals normalized between 0 and 1 and **varList** a vector containing the list of active ROIs.

#### 1.3 Signal segmentation and statistics extraction

Both the baseline correction and segmentation steps are implemented in the function *segWithBaseCorr* that is used as follows:

```
[meanFreq,realEvts,diffToBaseAll,varList] = segWithBaseCorr(...
dechNorm,dt,dtev,minDepth,maxEvThresh,Tev,cfDown,smoothParam);
```

where **dechNorm** is the matrix containing normalized signals, i.e the output from the function *extractIndivSignals*, **dt** is the acquisition time step (in seconds), **dtev** is the characteristic event half-time removed around peaks (in the present case we use $dtev = \Delta t_{ev} = 10$s), **minDepth** is the minimal peak depth for peak detection as described in the baseline correction step, **maxEvThresh** is the empirical max threshold for event detection (in our case we use 0.08), **Tev** is the ratio to noise (i.e. empirical standard deviation of the signal outside peaks) for event detection (4.5 in our case), **cfDown** is the ratio to **dtev** for the case were decay and increase in activation periods removal are different (here we use 1.5), and **smoothParam** is the smoothing spline parameter for the baseline fitting.
This function returns the following events features: **meanFreq** the mean event frequency for each ROI, **realEvts** a cell array containing all the event features for each ROI, organized as follows:

- column 1: beginning times $t_i$ of all detected events
- column 2: times of the main peak of all events
- column 3: times $t_e$ when all events end

- column 4: subsection number
- column 5: events amplitudes
- column 6: number $n_p$ of sub-events
- column 6: sub-events frequency.

The remaining outputs are **diffToBaseAll**: a matrix of the size of *dechNorm* containing the signals with their individual baseline correction, and **varList** a vector containing the list of ROIs that have at least one activation event detected.

#### 1.4 Detection of the co-activation paths

This step is implemented in two steps in the toolbox, first, the participating ROIs and activation order per global event are given by the function *globalEvtActiv*:

```
GloEvMat = globalEvtActiv(subperiodsTimes,realEvts,varList);
```

*globalEvtActiv* takes the inputs **subperiodsTimes** a vector that gives the timepoints of the subperiods segmentation, **realEvts** the struct containing all the calcium events information for all ROIs, obtained in the previous section with the function *segWithBaseCorr*, **varList** the list of active ROIs (over the total recording session), and returns the output **GloEvMat** a cell array containing the list of active ROIs and their peak time for each global event. Then, the co-activation paths are obtained with the function *activePaths* as follows:

```
[paths,area,numActiv,evPathLength,pathSteps,pathStepsTimes] = ...
activePaths(subperiodsTimes,GloEvMat,cellMap);
```

This function takes the inputs **subperiodsTimes** defined above, **GloEvMat** given by the previous function *globalEvtActiv*, and **cellMap** the map of the detected ROIs which is the output of *cellDetectFun* as described in section 1.1 of the main text. It returns **paths**, a cell array containing the activation paths of all global events in the form of a list of spatial coordinates (the ROI centroids) ordered along the activation path, **area** a vector of size $N$ containing the area in $\mu$m$^2$ of the convex hull of the active ROIs for each subperiod, **numActiv** a vector of size $N$ containing the number of active ROIs per subperiod, **evPathLength** a vector containing the total length in $\mu$m of the activation path for each subperiod. It also returns **pathsSteps**, a cell array of vectors containing the length (in $\mu$m) of each individual step of the path (each arrow in Fig.2E in the main text), and **pathStepsTimes** another cell array of vectors giving the time (in seconds) spent for each step.

#### 1.5 Construction of the connectivity graphs

The connectivity graphs and the information about the highly connected astrocytes are constructed using the function *activGraph*:

```
[netGraph, meanDeg, varDeg, numHighConnNodes, propHighConnNodes, conMat] = ...
activGraph(subperiodsTimes, GloEvMat, T_HC, orientation);
```

where **subperiodsTimes** is the list of timepoints for the segmentation into subperiods, **GloEvMat** the output of the function *globalEvtActiv*, $T_{HC}$ the high connectivity threshold, (an integer greater than 1), **orientation** a string stating whether the graph is oriented or not ('yes' or 'no'). It returns **netGraph** a MATLAB graph object, where the nodes are the ROI centroids and the edges weights are computed as explained in the main text, section 1.2, **meanDeg** the mean node degree (number of non zero edges) for each node = number of neighbors with direct connections to a specific node, averaged over all ROIs, **varDeg** the standard deviation among node degrees, **numHighConnNodes** the number of nodes with at least one edge of weight greater than $T_{HC}$, i.e. the number of ROIs that are part of the high connectivity subgraph, and **propHighConnNodes** the proportion of nodes with at least one edge of weight greater than $T_{HC}$, i.e. $propHighConnNodes = numHighConnNodes/nROIs$ where $nROIs$ is the total number of ROIs in the session (number of nodes of the entire graph). Finally, **conMat** is the connectivity matrix (directed or not according to the choice of input value **orientation**), in the case of non oriented graphs, the connectivity matrix is symmetric, in the case of oriented graphs it is not.

### 2 Supplementary results

#### 2.1 Individual activation statistics

From the segmented time-series (Fig. S1G), we extracted the individual activation statistics of each given ROI (with the function *segWithBaseCorr* described in subsection 1.3) and compared them between the two regions. Interestingly, we observed significant differences in the ROI activation at the individual scale. Specifically, the duration of activations (Fig. S2A) is longer and more variable in the CA1 ($4.48 \pm 3.83$ s, purple) compared to the CTX ($4.22 \pm 3.09$ s, yellow). On the contrary, event frequency (Fig. S2B-C) is higher in the CTX ($0.11 \pm 0.08$ Hz, yellow) than in the CA1 $0.08 \pm 0.07$ Hz. Accordingly, we found longer inter-events (Fig. S2D) in the CA1 ($26.94 \pm 31.88$ s) than in the CTX ($18.52 \pm 27.18$ s). Inter-events are not simply the inverse of event frequencies as the former is the quiescent time between the end of one event and the beginning of the next one (see grey lines in Fig. S2D), while the latter is the inverse of the duration between the main peaks of two consecutive events (black double arrow between the pink crosses in Fig. S2B-C). We also found higher amplitudes in the calcium events (Fig. S2E) in CA1 ($4.53 \pm 2.85$) compared to the CTX ($4.32 \pm 3.21$). Finally, the CA1 seems to have slightly more complex events ($1.33 \pm 0.02$ sub peaks per event) compared to the CTX ($1.25 \pm 0.02$, yellow), although this difference is not statistically significant.

#### 2.2 Stability analysis for graph reconstructions with respect to the number of calcium subsections

To guarantee that the network statistics are reaching a steady state as the number of subsections in the recordings $N$ increases, we used the following procedure: we ran the AstroNet-Construct full pipeline for increasing values of $N \in [\![0, 20]\!]$ and extracted the mean node degree of the ROIs (Fig. S3A) for all sessions of the two regions (CTX in yellow, CA1 in purple) for all values of $N$. Each curve shows the mean value of node degrees for a recording session, we then normalized the curves by the final value (for readability purposes). Indeed, all curves plateau at different values depending on the recording and the region, however the convergence always occurs quickly before $N = 20$ to a stable value, showing that the value $N = 20$ does not affect the results. Thus, we could reconstruct the graph properties, independently of the choice of $N$. Similarly, we obtain that the other properties measures: the variance in node degrees (Fig. S3B), the mean path length (Fig. S3C), as well as the number of hub astrocytes (Fig. S3D), the active area per subsection (Fig. S3E) and the number of active astrocytes per subsection (Fig. S3F) were all converging quickly to their final value as $N \in [0, 20]$ increases. To conclude, altogether these results show that the choice of the total number of subsections $N = 20$ was sufficient to reconstruct the main statistical properties of the network and that the convergence occurred before reaching this number.

### 3 Supplementary figures

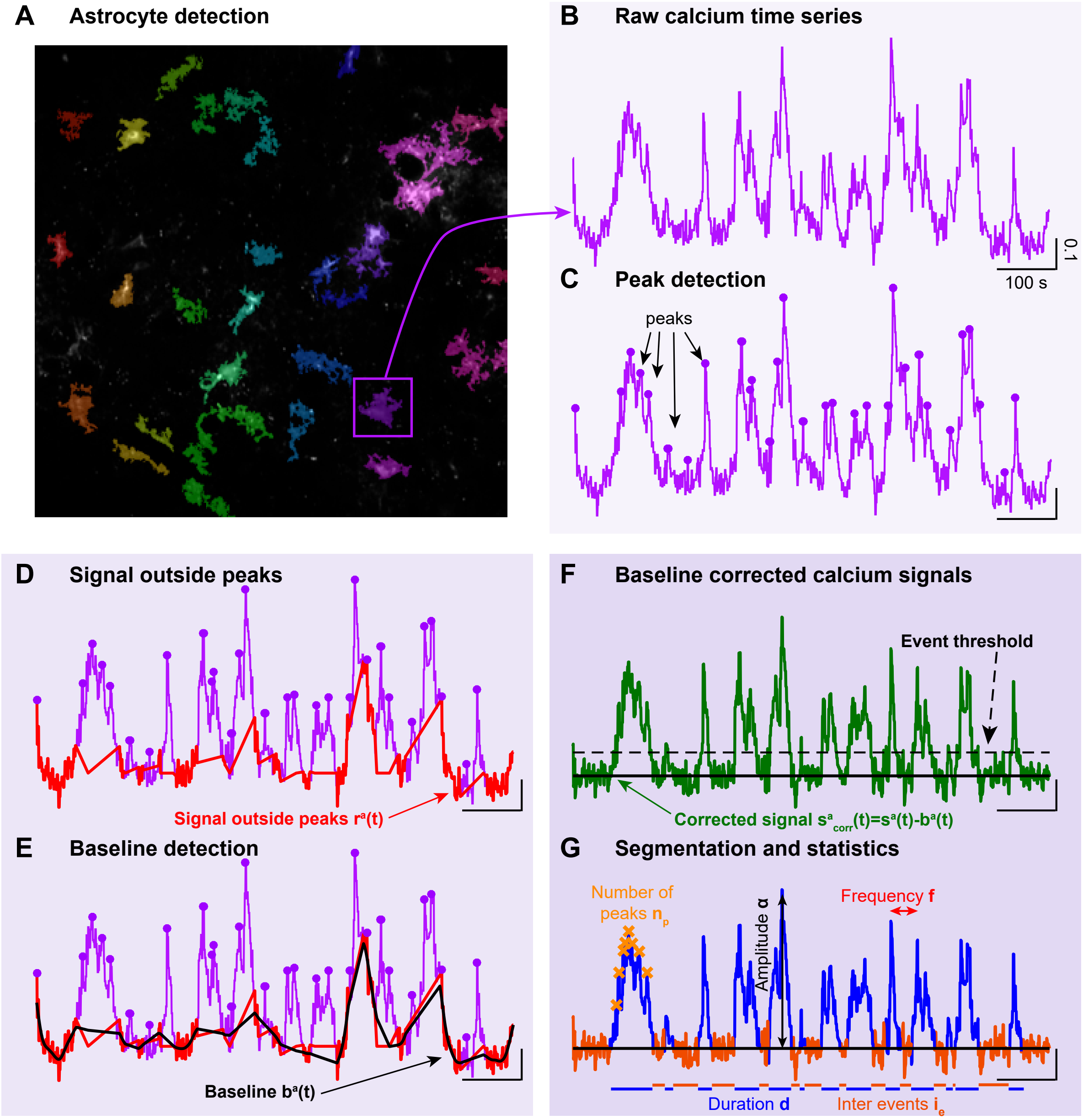


Figure S1: **Individual signal processing.** **A.** Detected ROIs (colored regions) from the summed image over the entire recording. **B.** Extraction of the average calcium time-series $s^a$ over all pixels of the ROI $a$ (purple, from the ROI in the square in panel A). **C.** Detection of the peak times (purple dots). **D.** Extraction of the signal outside of peaks (red) $r^a$ used for the baseline recovery. **E.** Reconstruction of the baseline (resting state) $b^a$ (black) using a smoothing spline on $r^a$. **F.** Corrected signal $s^a_{corr} = s^a - b^a$ (green) and event threshold detection based on the signal outside of peaks' standard deviation $E_T = 4std(r^a)$ (dotted black line). **G.** Signal segmentation and extraction of individual activation statistics for each ROI.

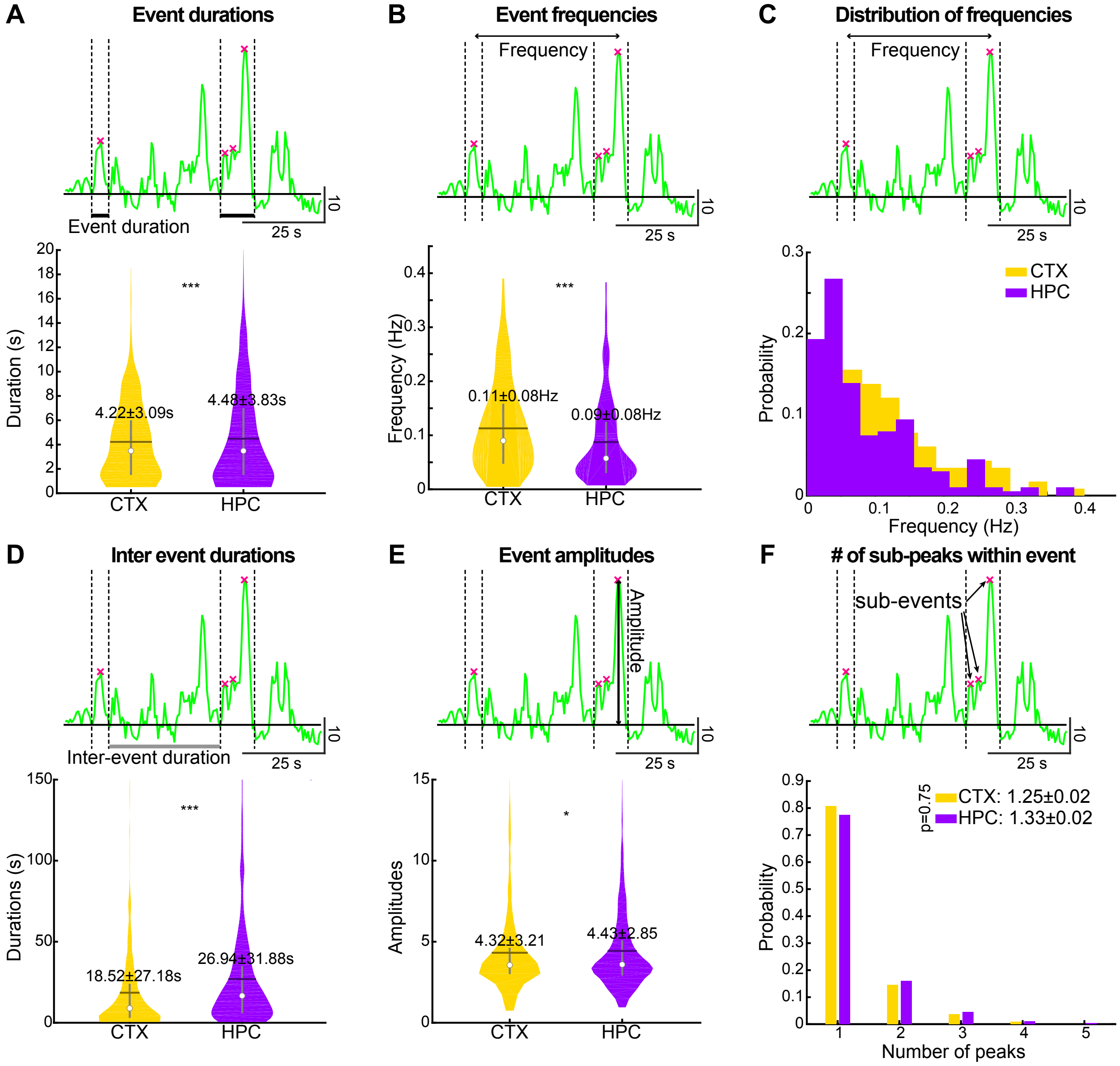


Figure S2: **Individual activations statistics.** **A.** Individual events durations, Upper: Single ROI time-serie showing the events duration extracted from the signal (thick black lines between dotted vertical lines), lower: violin plots for CTX (yellow) and CA1 (purple). **B.** Same for events frequencies. **C.** Distribution of events frequencies for the two regions. **D.** Same as A. and B. for inter-events durations (upper: thick grey lines). **E.** Same for individual events amplitudes. **.** **F.** Number of sub peaks (pink crosses in the upper plot) within each event.

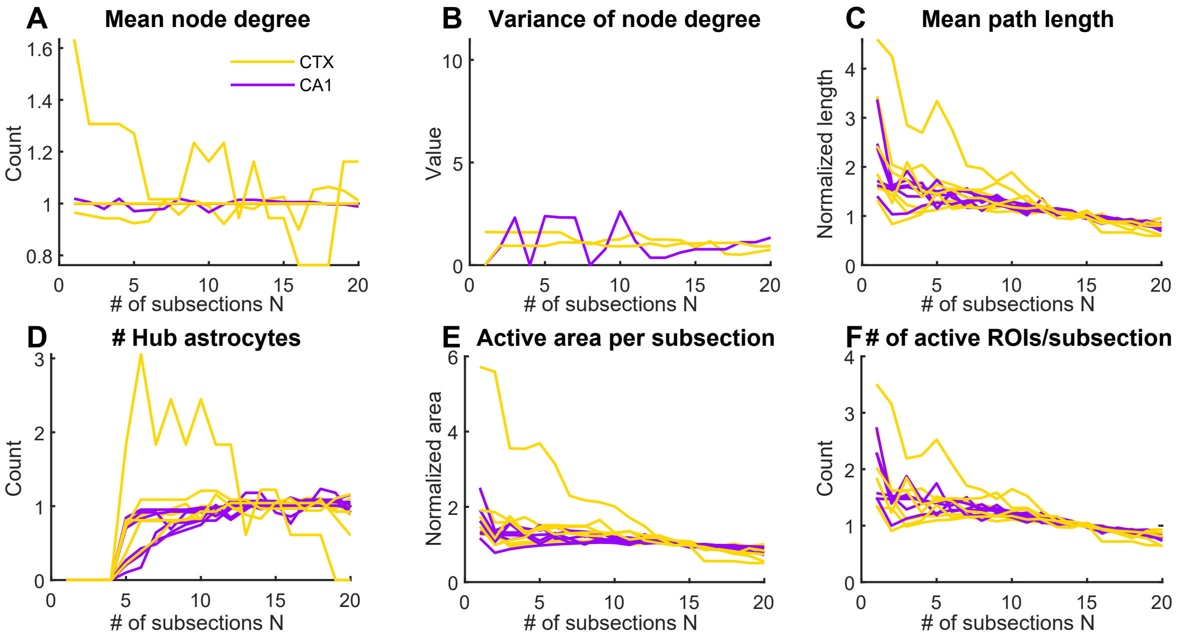


Figure S3: **Stability analysis of paths and networks reconstruction.** Convergence of various parameters (see below) vs number of subperiods $N \in [1, N_{max}]$, with $N_{max} = 20$. **A.** Mean node degree. **B.** Standard deviation of the node degree. **C.** Mean path length. **D.** Number of hub astrocytes. **E.** Area of the convex hull of the active astrocytes in each subperiod. **F.** Number of active astrocytes per subperiod. All values are normalized with respect to their final limits for readability. Each curve represents the results for one recording, color coded by brain region: CTX (yellow) and CA1 (purple).

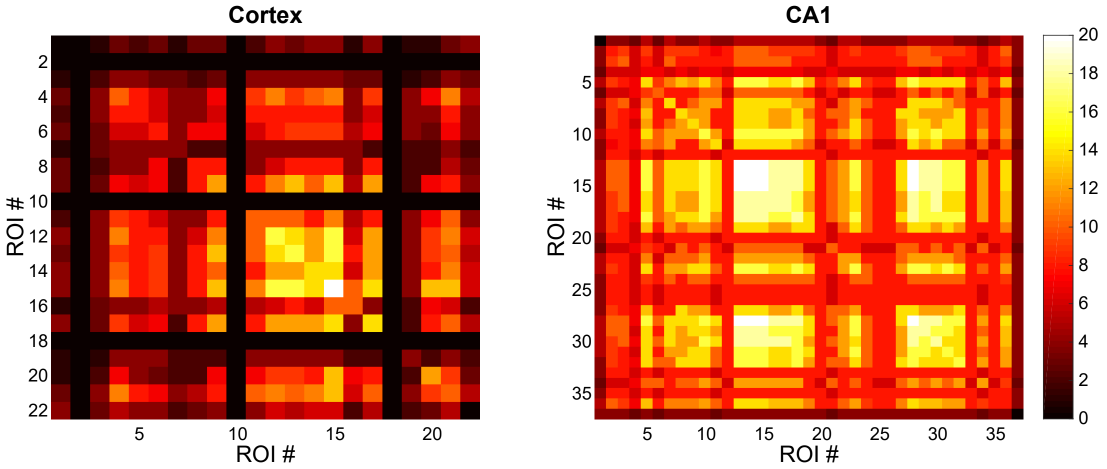


Figure S4: **Example of connectivity matrices** for one representative recording in each region. Left: CTX, right: CA1, colorbar indicates the strength of connection

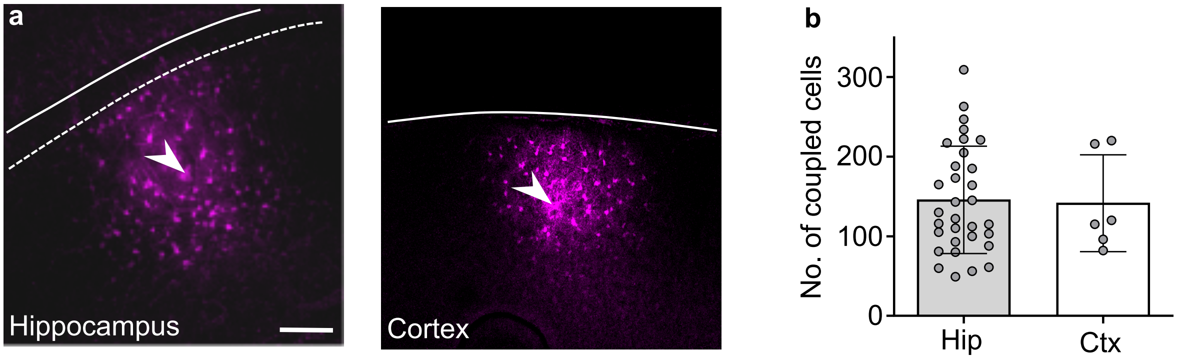


Figure S5: **Network reconstructed from passive diffusion through gap junction.** Similar size of gap-junction mediated astroglial networks in the hippocampus and cortex. (a) Representative images of astroglial networks in the hippocampus (CA1 area) or cortex, assessed by intercellular coupling in astrocytes patched with a pipette filled with biocytin. The white dotted lines show the borders of the hippocampal pyramidal cell layer and cortex, respectively. The white arrow heads indicate the injection site. (b) Quantification of the size of the astroglial networks in the hippocampus (n=32) and in the cortex (n=6).

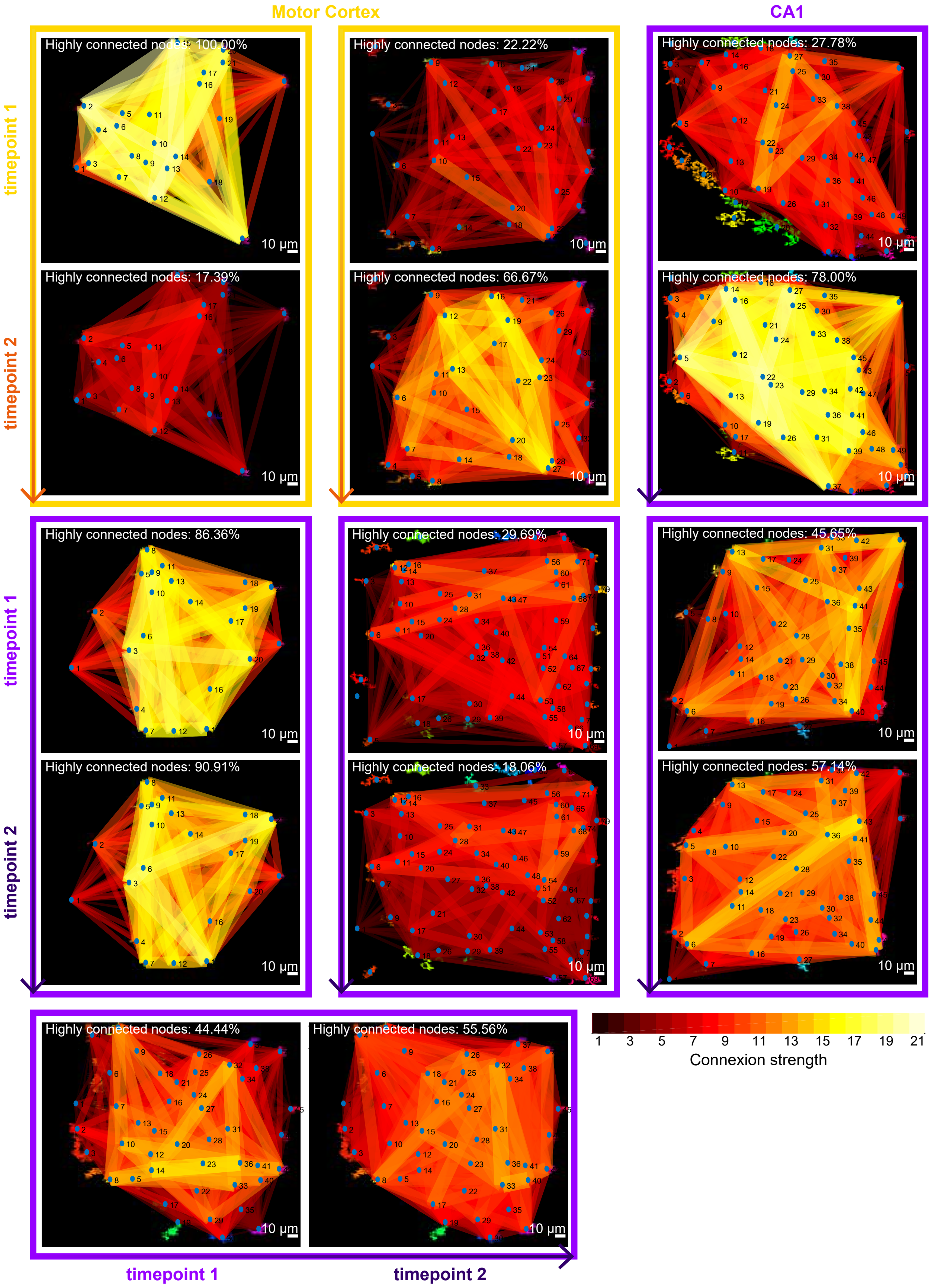


Figure S6: **Time-lapse analysis of graph stability.** Examples of reconstructed graphs at consecutive timepoints 1 (upper) and 2 (lower, 5 min later) in the same slice, for two CTX slices (yellow) and 5 CA1 slices.

### 4 Comparison to other calcium analysis softwares

#### 4.1 Overview

The main goal of the present manuscript is to present a computational approach to extract ROI (of astrocytes) and to evaluate their functional connectivity at the local network scale. Furthermore, this method allows us to quantify and compare the functional network between different brain regions (here CTX and CA1). Since there exist many different tools for calcium analysis, we provide here a quick overview:

- **GECI-quant**: is optimized for analyzing calcium imaging data, particularly GCaMP-based data for neuronal calcium activity. It processes the data to extract metrics like amplitude, frequency, and duration of calcium transients. This tool is primarily neuron-focused and does not have specialized algorithms for analyzing astrocyte networks or connectivity. While it can detect calcium signals, estimating astrocyte connectivity would require custom adaptations.

- **Suite2P** is designed for fast and efficient analysis of large calcium imaging datasets. It performs motion correction, ROI extraction, and signal extraction from calcium imaging data, primarily in neurons. Suite2P uses a combination of sparse data representation and fast clustering to separate neuronal signals. Suite2P is neuron-centric and does not inherently support astrocyte-specific connectivity analysis. While it could be adapted for astrocyte data, its algorithms are not designed to address the unique characteristics of astrocytic calcium signaling, such as the slower and more global nature of their calcium waves. The lack of astrocyte-specific algorithms limits its effectiveness for estimating astrocyte connectivity.

- **LC_Pro** is specifically tailored for analyzing calcium signals in glial cells, particularly astrocytes. It emphasizes the detection of slower calcium transients typical of astrocyte activity and includes tools for analyzing spatiotemporal patterns in glial cell calcium signals. while LC_Pro allows for the analysis of calcium dynamics within individual astrocytes, it is not explicitly designed to infer or model connectivity between astrocytes. Analyzing network-level interactions would require additional analytical layers to estimate connectivity based on shared or synchronized calcium signals.

- **CaSCaDE** is primarily developed for neuronal calcium data analysis and spike inference, helping to decode action potentials from calcium transients. The software emphasizes extracting neuronal firing information from calcium signals. CaSCaDE is primarily focused on neurons and the inference of spikes, it is not suitable for analyzing astrocytic activity or connectivity. The algorithms in CaSCaDE are optimized for neuron spiking rather than the continuous, slower dynamics seen in astrocytes.

- **CaImAn** is a versatile and powerful tool for analyzing calcium imaging data, featuring motion correction, source extraction, and deconvolution algorithms. Its advanced techniques allow for highly accurate calcium signal extraction and temporal deconvolution. CaImAn could be adapted for use in both neuronal and astrocyte data, making it one of the more flexible tools. Its strong signal extraction and motion correction algorithms allow it to handle astrocyte calcium imaging data. However, while CaImAn can process the data efficiently, it lacks built-in modules similar to AstroNet specifically for estimating astrocyte connectivity. With custom post-processing steps or by integrating connectivity analysis methods, it could be used to infer astrocyte networks, but this would require significant adaptation and coding.

- **STARDUST** is designed for the spatial and temporal analysis of calcium signaling patterns in glial cells (including astrocytes) and neurons. It focuses on detecting patterns in calcium dynamics and is particularly suited for analyzing mixed populations of glial and neuronal cells. STARDUST offers more promise in terms of astrocyte connectivity analysis. By detecting and analyzing calcium signaling patterns in astrocytes, it could be used to infer astrocyte connectivities similar to AstroNet, especially when paired with its ability to track calcium signals across time and space. This tool is better suited than the previous ones for assessing astrocyte interactions based on calcium wave propagation and correlated activity. However, it does not recover this reconstruction as presented by AstroNet.

#### 4.2 Detailed comparison with AQuA2

AQuA2 is a new, unpublished version of AQuA, posted on BioRxiv in June 2024. Unlike the original AQuA, AQuA2 now incorporates ROI segmentation. By improving the event segmentation process, AQuA2 merges events by considering delayed calcium dynamics and spatial overlap, forming distinct Consensus Functional Units (CFUs). We compared AstroNet's segmentation pipeline with AQuA2 using the hippocampus and motor cortex datasets. The comparison employed Dice scores, confusion matrices, and accuracy metrics. Overall, AQuA2 achieved a slightly better F1 score, indicating improved predictions, despite having similar accuracy to AstroNet. Both algorithms successfully detect astrocytes (as seen in the proportion of false negatives) but tend to overestimate astrocyte counts (reflected in the false positives). Filters can be applied in both software to fine-tune astrocyte detection. AQuA2 also provides additional features, such as propagation maps and event-based statistics. However, AstroNet significantly outperforms AQuA2 in speed. Most importantly, AstroNet offers a complete network reconstruction method based on graphs and activation paths, a capability entirely absent in AQuA2. As previously mentioned, the two software serve different purposes, with AstroNet delivering critical complementary information for quantifying local network activity.

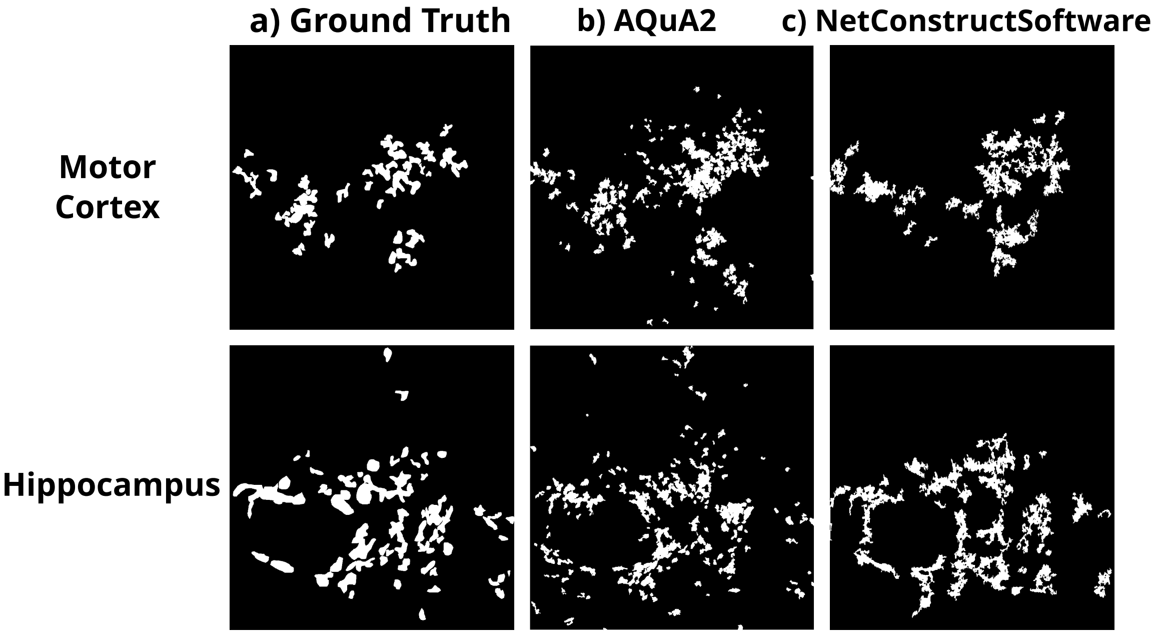


Figure S7: Comparison of the binary segmentation of astrocytes with AQuA2 vs AstroNet in the motor cortex and hippocampus datasets.

| Dice (or F1 score) | AstroNet | AQuA2 |
|---|---|---|
| CTX | 0.45 | 0.54 |
| CA1 | 0.47 | 0.53 |
| **Accuracy** | **AstroNet** | **AQuA2** |
| CTX | 0.941 | 0.943 |
| CA1 | 0.906 | 0.921 |

Table 1: **Comparison of AstroNet and AQuA2**

| CTX | | CA1 | |
|---|---|---|---|
| 0.41 | 0.59 | 0.46 | 0.54 |
| 0.03 | 0.97 | 0.05 | 0.95 |

Table 2: **Confusion matrix of AstroNet**

| CTX | | CA1 | |
|---|---|---|---|
| 0.45 | 0.55 | 0.56 | 0.44 |
| 0.02 | 0.98 | 0.05 | 0.95 |

Table 3: **Confusion matrix of AQuA2**
